# Regional Hypothalamic Responses to Light Vary Across Developmental Stages but Remain Stable with Time of Day

**DOI:** 10.64898/2026.01.05.697645

**Authors:** Roya Sharifpour, Islay Campbell, Fermin Balda Aizpurua, Ilenia Paparella, Elise Beckers, Nasrin Mortazavi, Fabienne Collette, Christophe Phillips, Puneet Talwar, Laurent Lamalle, Mikhail Zubkov, Gilles Vandewalle

**Author notes:** Corresponding authors: Gilles Vandewalle, GIGA-Cyclotron Research Centre-Human Imaging, Bâtiment B30, 8 Allée du Six Août, University of Liège-Sart Tilman, 4000 Liège, Belgium. Roya Sharifpour, Department of Translational Neuroimaging, Hôpital Erasme, Hôpital Universitaire de Bruxelles, Université libre de Bruxelles, Brussels, Belgium. Department of Translational Neuroimaging, Hôpital Erasme, Hôpital Universitaire de Bruxelles, Université libre de Bruxelles, Brussels, Belgium (Roya Sharifpour). Sir Jules Thorn Sleep and Circadian Neuroscience Institute, Nuffield Department of Clinical Neurosciences, University of Oxford, Oxford, UK. Shared first authorship (Roya Sharifpour, Islay Campbell). Joint senior authorship (Mikhail Zubkov, Gilles Vandewalle).

## Abstract

**Background:** Light plays a significant role in regulating various non-visual biological processes, such as stimulating alertness and cognition. However, the precise subcortical neural pathways are not fully established, including within the hypothalamus. In particular, how the hypothalamus processes are modulated by time-of-day and developmental stages, remains poorly understood. **Methods:** In this study, we used 7 Tesla functional magnetic resonance imaging to examine *in vivo* the impact of different light illuminance (0.16, 37, 92, and 190 melanopic equivalent daylight illuminance: mel-EDI lux) on the activity of the hypothalamus of healthy young adults (N=33; 20 women; 24.3 ±3.2y) and adolescents (N=16; 5 women; 16.8 ±1.1y) while they completed an auditory executive task, in the morning or in the evening. **Results:** Performance to the task improved with increasing illuminance irrespective of time of day and age group. When focusing on time-of-day differences in young adults, we found that the regional impact of light illuminance on the activity of the hypothalamus was consistent between the morning and the evening, with the posterior and anterior hypothalamus, respectively, showing increased and decreased activity with increasing illuminance. When focusing on developmental stages differences, during the evening session only, we found similar regional patterns in adolescents and young adult. The magnitude of the response at the highest illuminance was, however, larger in adolescents, with a larger deactivation of the superior-anterior and inferior-tubular hypothalamus. **Conclusions:** These findings reveal a complex and non-uniform impact of light on hypothalamus activity and provide novel insights into how light influences vary with developmental stages.

## Background

Light exerts multiple impacts on human physiology extending beyond vision through non-image-forming (NIF) effects that regulate key biological processes, including circadian rhythmicity, sleep-wake regulation, alertness, mood and cognitive function ^1^. These NIF effects are largely mediated by intrinsically photosensitive retinal ganglion cells (ipRGCs) which are maximally sensitive to short-wavelength (blue) light (∼480nm) ^2^. IpRGCs combine their intrinsic photosensitivity with the signals of rods and cones to relay light signals to a broad range of brain regions ^1^.

A critical target of ipRGC signaling is the hypothalamus. Several distinct hypothalamic nuclei receive ipRGC inputs, including over the anterior part, with the suprachiasmatic nucleus (SCN), consisting of the master circadian pacemaker, and the preoptic area nuclei (POA), which promotes sleep, as well as over the lateral and posterior parts, with the lateral hypothalamus (LH), containing both wake- and sleep-promoting neurons ^3–7^. These nuclei are considered to play key roles in modulating light’s effects on arousal, alertness, and cognition. Much of our understanding of the impact of light on hypothalamus nuclei comes, however, from animal studies ^1,2^, where neural circuits can be invasively studied and manipulated. Translating these findings to humans is not straightforward, due to differences in anatomy, physiology, and complexity of the human cortex ^8^. The activity of the anterior part of the hypothalamus, which largely encompasses the SCN, was recently reported to decrease with increasing illuminance during various cognitive tasks ^9^ as well as in the absence of a cognitive task ^10^ using functional magnetic resonance imaging (fMRI). In contrast, the posterior part, including part of the LH, showed the opposite pattern during cognitive tasks, with increasing activity at higher illuminance ^9^.

The NIF impact of light varies both with environmental and individual factors ^1^. Among environmental factors, time-of-day has an essential impact on the entrainment of the circadian clock ^1^ and has been reported to affect the stimulating effect of light on human alertness and cognition ^11,12^. In terms of individual factors, age affects light impact on circadian entrainment, alertness and cognition. Most of the evidence pertain, however, to older age, while adolescents may also exhibit different sensitivity and neural responses to light ^13^. Whether time-of-day and specificities of teenagers can be detected at the level of the hypothalamus is not known.

In this study, we took advantage of high-resolution 7-Tesla fMRI to investigate how the regional impact of light illuminance changed within the hypothalamus with time-of-day and between adolescence and early adulthood. Specifically, we sought to determine whether the hypothalamic response of young adults (19-30 years) to light varied from the morning to the evening. We focused on the executive functions, as it has been repeatedly used successfully to uncover how light affect cognitive brain function including at different times of day ^12^. Because of the current concerns about their evening light exposure ^14^, we further included a group of late teenagers (15–18 years) who completed the protocol only in the evening. We hypothesized that in young adults, NIF effects would be larger in the morning, while in the evening, NIF responses would be larger in adolescents compared to young adults.

## Methods

This cross-sectional research is part of a broader investigation that has led to several publications using part of the adult participants included in the present paper ^9,15–17^. All procedures and analyses are the same as those in ^9^, except for participant groups and analyses pertaining to group comparisons. The Ethics Committee of the University of Liège approved the study. All the participants provided written informed consent and were financially compensated for their participation.

### Participants

This study involved 55 healthy participants aged 15-30 years (22.0 ± 4.6 years; 28 women) recruited between February 2021 and September 2023. The study consists of between-group comparisons including 3 non-overlapping groups of participants: 20 young adults completed the protocol in the morning (24.2 ± 2.5 years; 13 women; 19 of which were included in ^9^); 17 young adults completed the protocol in the evening (25.2 ± 4.0 years; 10 women; 6 of which were included in ^9^ but with their data collected in the morning); 18 adolescents completed the protocol in the evening (16.7 ± 1.1 years; 5 women). Due to practical constraints related to school schedules, morning data collection was deemed too difficult for the adolescent group in the context of our experiment.

The exclusion criteria for all groups included a Body Mass Index (BMI)>28, recent psychiatric history, severe trauma, sleep disorders, addiction, chronic medication use, smoking, excessive alcohol consumption (>14 units/week), and high caffeine intake (>4 cups/day). Participants were also excluded if they had chronic night shift work in the past year, transmeridian travel in the past two months, or any history of ophthalmic disorders. Participants were included if they scored below 18 on the 21-item Beck Anxiety Inventory (BAI) (indicating mild anxiety or less) ^18^, below 14 on the Beck Depression Inventory-II (BDI-II) (indicating mild depression or less) ^19^, below 8 on the Pittsburgh Sleep Quality Index (PSQI) (indicating good sleep quality) ^20^, and below 12 on the Epworth Sleepiness Scale (ESS) (indicating normal daytime sleepiness) ^21^. Participants also completed questionnaires assessing chronotype (Horne-Östberg)^22^ and seasonal mood variation (Seasonal Pattern Assessment Questionnaire - SPAQ)^23^, but these questionnaires were not used for participant inclusion. The demographic details of the participants are summarized in Table 5.

### Study Design and Procedure

To prevent excessive sleep deprivation while maintaining realistic entrained conditions, participants were instructed to follow a loose sleep‒wake schedule for seven days prior to their laboratory visit, with bed and wake times restricted to within ±1 hour of their habitual schedule. Compliance was verified using actigraphy (AX3 accelerometer, Axivity, United Kingdom) and daily sleep diaries. Additionally, participants were requested to refrain from all caffeine- and alcohol-containing beverages, as well as extreme physical activity, for 3 days before participating in the fMRI acquisition. The timing of the experiments varied based on group assignment: the morning session started approximately 2.5 hours after the participants’ habitual wake-up time, and the evening session started approximately 1 hour before their habitual bedtime. Upon arrival, all participants underwent a light adaptation protocol to standardize recent light exposure before the fMRI scan. This involved 5 minutes of exposure to relative bright polychromatic white light (∼1000 lux), followed by 45 minutes of ambient dim light (<10 lux). During this period, the participants received instructions and completed practice trials of the cognitive task (n-back) on a laptop (**Figure 1-A**).

**Figure 1:**
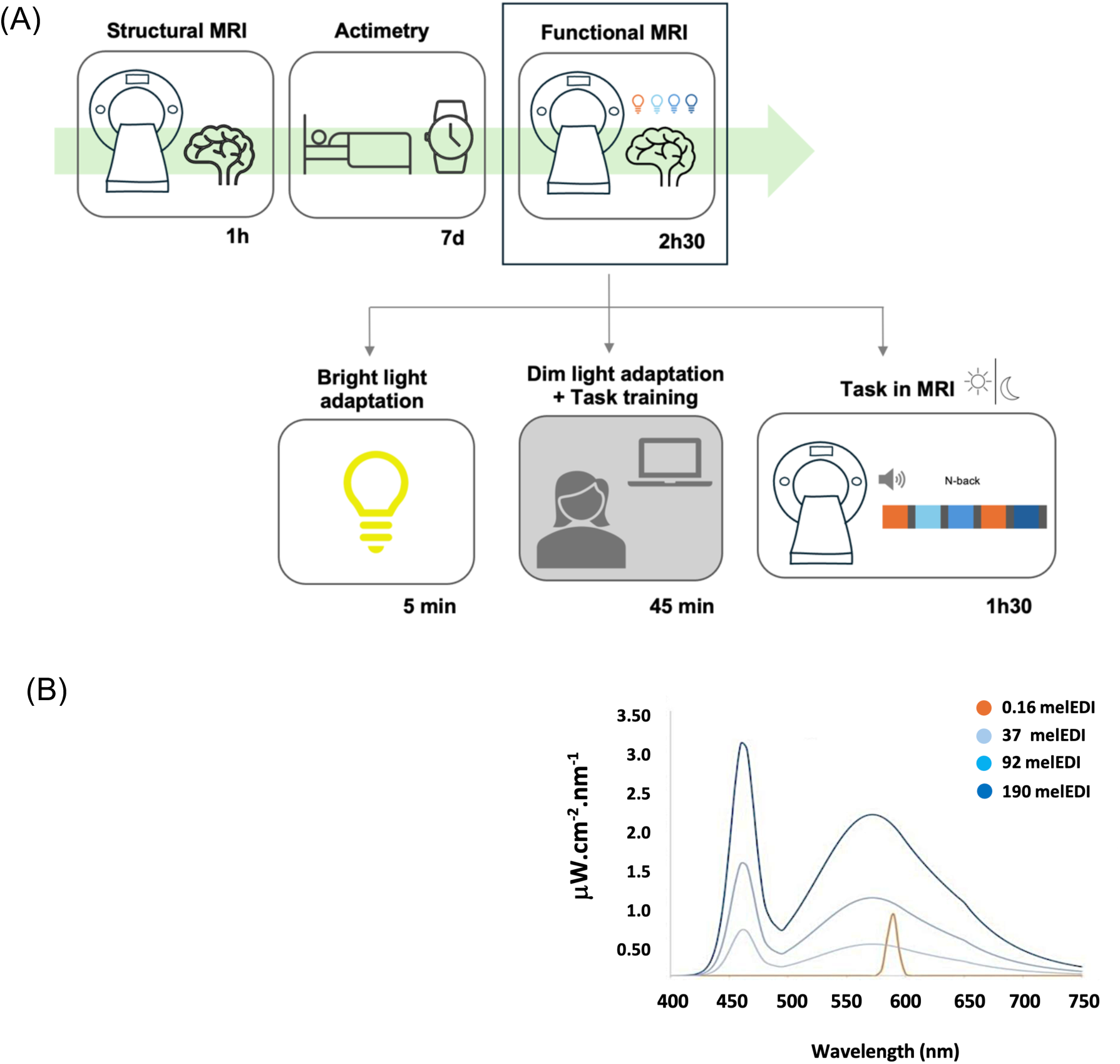
Experimental protocol: **A)** Participants first underwent a structural MRI session, followed by 7 days of monitored regular sleep-wake schedule and an fMRI session. To standardize recent light history, participants were exposed to bright white light (1000 lux) for 5 minutes and were maintained in in dim-light for 45-minute (<10 lux) during which they practiced the fMRI auditory tasks. In the MRI apparatus, participants performed an N-back working memory task under alternating blocks of polychromatic blue-enriched and monochromatic orange light, separated with darkness intervals. The duration of each step is indicated below the respective timeline elements. **B)** Spectral profiles of the four light conditions used during the N-back task (detailed light characteristics can be found in Table S1).

Inside the scanner, an auditory letter variant of the n-back task, a well-established measure of working memory ^24^, was presented to the participants. In the letter detection 0-back condition, participants were instructed to respond whenever the current auditory stimulus matched a predefined target letter (the letter “K”). This condition served as control for baseline brain activity changes. In the 2-back condition, participants were required to continuously update and maintain relevant information in working memory. They had to identify whether the current auditory stimulus matched the auditory stimulus presented two items earlier. The task was delivered in a block design, alternating in a pseudo-random manner (ensuring the spread of each task type over the entire recording) between blocks of 0-back and 2-back tasks (38 blocks in total consisting of 19 blocks of 0-back and 19 blocks of 2-back), yielding a total task duration of approximately 28 minutes. The participants responded using an MRI-compatible keypad held in their dominant hand. Auditory stimuli were delivered through noise-cancelling, MRI-compatible headphones to ensure clear auditory perception despite the MRI noise. The n-back task was followed by an emotional and an attentional task, which will not be discussed in the present paper.

Participants were alternately maintained in darkness (<0.01 lux) or exposed to blue-enriched cool polychromatic light (6500K) at varying illuminance levels (37, 92, and 190 mel-EDI lux) and a control monochromatic orange light (590 nm; 0.16 mel-EDI lux), to which ipRGCs are nearly insensitive (**Figure 1-B**). Orange light was introduced as a control for visual stimulation for potential secondary whole-brain analyses. For the present region of interest analyses, we discarded color differences between the light conditions and only considered illuminance as indexed by mel-EDI lux. This constitutes a limitation of our study, as it does not, for instance, allow us to attribute the findings to a particular photoreceptor class.

There were 7 task blocks (each lasting ∼30 seconds) for each light condition and 6 blocks for the darkness condition, totaling 34 light blocks across the experiment. The N0 and N2 task conditions were presented pseudo-randomly within these blocks, with each task occurring at least three times under each light condition. Periods of darkness (∼10 s, <0.01 lux) without tasks separated blocks with different light conditions. In some instances, two same task blocks occurred consecutively under the same light condition without an intervening darkness period, effectively forming a continuous ∼60-second light segment. In some cases, two same task blocks with the same light condition were followed by a short (∼10 s) rest period under the same light condition, resulting in a ∼70-second light segment. During all the fMRI sessions, an eye-tracking device (SR-Research, Canada). confirmed that the participants kept their eyes open during the scan. The entire experiment was designed using OpenSesame^25^.

### MRI Data Acquisition

Structural and functional MRI data were obtained using a MAGNETOM Terra 7 Tesla (7T) MRI system (Siemens Healthcare, Erlangen, Germany) equipped with a 32-channel receiver and 1-channel transmit head coil (Nova Medical, Mam USA). To minimize dielectric artifacts, dielectric pads (Multiwave Imaging, Marseille, France) were positioned between the participants’ heads and the receiver coil. A multi-band Gradient-Recalled Echo - Echo-Planar Imaging (GRE-EPI) sequence was used to collect multislice T2*-weighted fMRI images, with the axial slice orientation and parameters set as follows: TR = 2340 ms, TE = 24 ms, FA = 90°, no interslice gap, field of view (FoV) = 224 mm × 224 mm, matrix size = 160 × 160 × 86, and voxel size = (1.4 × 1.4 × 1.4) mm³. The first three scans were discarded to reduce saturation effects. To control for physiological noise in the fMRI data, participants’ pulse and respiration were recorded using a pulse oximeter and a breathing belt (Siemens Healthineers, Erlangen, Germany). Following the fMRI scan, a 2D gradient echo (GRE) field mapping sequence was used to evaluate B0 magnetic field inhomogeneity. The acquisition parameters for the field mapping were as follows: TR = 5.2 ms, TEs = 2.26 ms and 3.28 ms, FA = 15°, bandwidth = 737 Hz/pixel, matrix size = 96 × 128, 96 axial slices, and voxel size = (2×2×2) mm³, with a total acquisition time of 1:38 minutes.

For anatomical reference, a high-resolution T1-weighted image was captured using a Magnetization-Prepared with 2 Rapid Gradient Echoes (MP2RAGE) sequence, with parameters including TR = 4300 ms, TE = 1.98 ms, FA = 5°/6°, TI = 940 ms/2830 ms, bandwidth = 240 Hz, matrix size = 256 × 256 × 224, acceleration factor = 3, and voxel size = 0.75 × 0.75 × 0.75 mm³.

### Data Preprocessing

Structural MP2RAGE images underwent denoising using Statistical Parametric Mapping (SPM12) with a method detailed in ^26^. These denoised images were subsequently automatically reoriented using SPM and corrected for intensity bias caused by field inhomogeneity using the bias correction method within SPM segmentation. Brain extraction was performed on the denoised, reoriented, and bias-corrected images to avoid potential coregistration issues arising from the use of dielectric pads. This process was carried out using SynthStrip ^27^. The fMRI time series preprocessing included 1) realignment and unwarping to correct for head motion and field inhomogeneities; 2) brain extraction, using SynthStrip; and 3) smoothing, which was applied with a Gaussian kernel (3 mm full width at half maximum) to improve the signal-to-noise ratio.

### Statistical Analyses

For first-level analysis, each subject’s data was analyzed in their native space to minimize errors from coregistration. The whole-brain univariate analyses consisted of a general linear model (GLM) computed with SPM12. Task and light blocks were modeled as block functions and convolved with the canonical hemodynamic response function. The fMRI time series was high-pass filtered (cutoff = 256 seconds) to remove low-frequency drifts. Movement and physiological parameters (heart rate and respiration), which were calculated using the PhysIO Toolbox (Translational Neuromodeling Unit, ETH Zurich, Switzerland), were included as covariates of no interest ^28^.

Two separate analyses were completed. The primary analysis aimed to assess whether brain responses during the task were influenced by overall changes in illuminance levels. For each task type, the task block regressors were accompanied by a single parametric modulation regressor reflecting the light melanopic illuminance levels (0, 0.16, 37, 92, and 190 mel-EDI lux). The contrasts of interest focused on the main effects of this parametric modulation. In the follow-up post hoc analysis, we evaluated responses to stimuli under each light condition. Separate regressors represented each task level under each light condition (0, 0.16, 37, 92, and 190 mel-EDI lux). The contrasts of interest consisted of the main effects of each regressor.

Using a deep learning approach implemented in FreeSurfer ^29^, we segmented the hypothalamus into five subregions: inferior-anterior, superior-anterior, posterior, inferior-tubular, and superior-tubular (**Figure 2-A**). We then utilized the output masks from this segmentation to extract regression betas (activity estimates) for each hypothalamic subregion using the REX Toolbox ^30^. The activity estimates were then averaged (means) within each subpart and across both hemispheres. In the main analyses, this resulted in one activity estimate for each task stimulus type and each hypothalamic subregion (totaling 10 per individual). In the subsequent analyses, we obtained five activity estimates for each task stimulus type for each subregion (total of 50 per individual).

**Figure 2:**
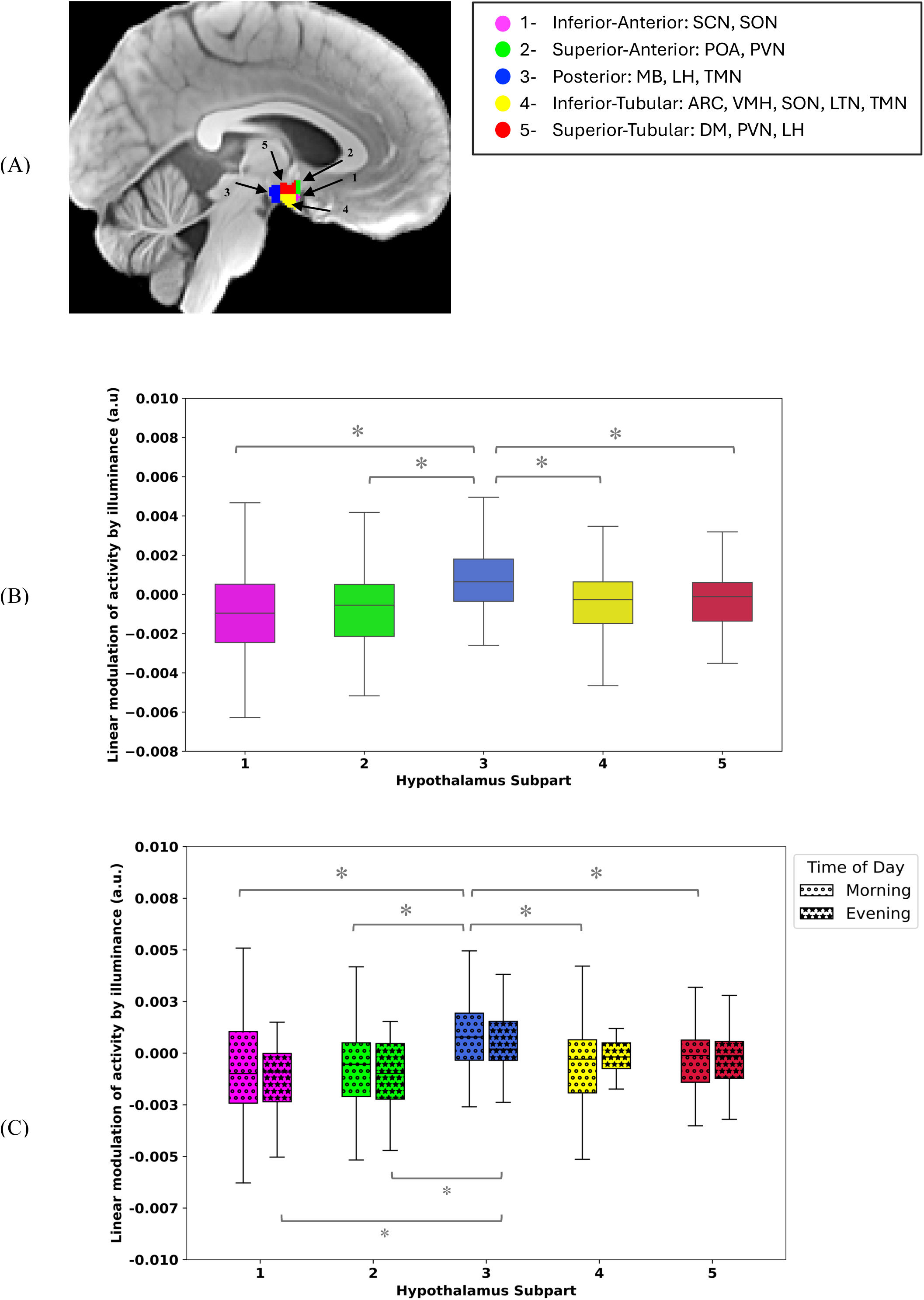
Regional hypothalamus response to light in young adults in the morning and evening groups. **(A)** Hypothalamus parcellation into five subparts encompassing the nuclei mentioned in the right inset. ARC: arcuate nucleus; DMH: dorsomedial nucleus; LH: lateral hypothalamus; LTN: lateral tubular nucleus; MB: mamillary body; POA: preoptic area; PVN: paraventricular nucleus; PNH: posterior nucleus of the hypothalamus; SCN: suprachiasmatic nucleus; SON: supraoptic nucleus; TMN: tuberomammillary nucleus; VMN: ventromedial nucleus. **(B)** Linear modulation of activity by illuminance (arbitrary unit – a.u.; median (horizontal line), interquartile range (IQR; boxed region), and whiskers extending to Q3 + 1.5 × IQR and Q1 – 1.5 × IQR) of each hypothalamus subpart in response to illuminance variation (Morning and evening subject together; N=33) showing increased impact of light in subpart three (posterior) relative to the other subparts (main effect of hypothalamus subpart: p < 0.0001). **(C)** Group comparison of linear modulation of activity by illuminance (a.u. ; median (horizontal line), interquartile range (IQR; boxed region), and whiskers extending to Q3 + 1.5 × IQR and Q1 – 1.5 × IQR) of each hypothalamus subpart in response to illuminance variation in the morning (N=18) vs. in the evening (N=15) (main effect of time of day: p = 0.84; interaction of hypothalamus subpart and time of day: p = 0.59) The linear modulation of activity by illuminance of the N0 and N2 tasks are averaged (task are presented separately in **Suppl. Figure S1**). Asterisks (*) denote significant differences (p < 0.05).

Statistical analyses of hypothalamic activity estimates were conducted using SAS 9.4 (SAS Institute, NC, USA). The analyses employed generalized linear mixed models (GLMMs), incorporating the subject as a random factor for both the intercept and slope, and were adjusted for the distribution of the dependent variable. Direct post hoc tests were adjusted for multiple comparisons using the Tukey method. Subsequent detailed analyses were treated as post hoc and did not undergo multiple comparison corrections (p < 0.05). To identify outliers in the datasets, Cook’s distance greater than 1 was utilized for exclusion. This process revealed two outliers in the activity estimates for the evening adult, morning adult, and adolescent groups. Semi-partial R^2^ (R^2^β*) values were computed to estimate the effect sizes of significant fixed effects and statistical trends in all GLMMs ^31^.

The primary analyses focused on activity estimates modulated by light illuminance as the dependent variable. The hypothalamic subpart and task stimulus type (2-back/0-back) were included as repeated measures (with compound symmetry structure), with age, sex, BMI, and season serving as covariates, along with an interaction term between hypothalamic subpart and time of day/age group. The subsequent set of post hoc GLMM analyses examined the activity estimates of the hypothalamic subparts as the dependent variable and the hypothalamic subpart, stimulus type, and illuminance levels (0, 0.16, 37, 92, and 190 mel-EDI lux) as repeated measures (with compound symmetry structure), with age, sex, BMI, and season as covariates, along with interaction terms between illuminance, hypothalamic subpart and time of day/age group.

A separate set of GLMMs was employed to assess: (1) whether illuminance levels influenced task performance, (2) whether time of day and/or age group affect performance, and (3) whether performance was associated with activity of any hypothalamic subregions. To address the first two questions two models designed: in one model, the task performance of all adult participants was analyzed. Task accuracy under different light conditions served as the dependent variable, with random effects for each subject (intercept and slope) along with illuminance levels (0, 0.16, 37, 92, and 190 mel-EDI lux) as repeated measure (autoregressive (1) correlation). The independent variables were illuminance level, time of day, and their interaction. Sex, BMI, age and season were included as covariates. A second model focused on participants scanned in the evening. Again, task accuracy under different light conditions was the dependent variable, and subject-specific intercepts and slopes were modeled as random effects along with illuminance level as repeated measure (autoregressive (1) correlation). This model included illuminance level, age group, and their interaction as independent variables. Sex and BMI were treated as covariates. As the final step, activity estimates of each hypothalamic subpart under each light condition added separately to the previous models to investigate whether there is a correlation between task performance and activity of any hypothalamic subpart.

Although optimal sensitivity and power estimation methods for generalized linear mixed models (GLMMs) are still being refined ^32^, we conducted a prior sensitivity analysis to approximate the smallest effect size that could be detected in our primary analyses, given our sample sizes. Using G*Power 3 (version 3.1.9.4; ^33^), and assuming a power of 0.8 and an alpha level of 0.05, our sample sizes (N=31, i.e. the smallest sample size of our analyses , when comparing young adults: N=15 and adolescents in the evening: N=16) provided sufficient power to detect large effect sizes (r > 0.47; two-tailed; absolute values; R² > 0.22, R² CI: 0.13–0.71). This analysis was framed within a multiple linear regression model including one predictor of interest (group) and at least four covariates (time of day/age group, task, sex, and BMI).

## Results

Forty-nine healthy participants completed the protocol and were included in the analyses: 18 young adults scanned in the morning (12 women, 24.0±2.4 y), 15 young adults scanned in the evening (8 women, 24.6±3.8 y), to assess time of day variations in young adults, and 16 adolescents scanned in the evening (5 women, 16.8±1.1 y) to assess age differences in the evening (**Table 1**). Their hypothalamus was parcellated into five subparts, inferior-anterior, superior-anterior, posterior, inferior-tubular and superior-tubular (**Figure 2-A**), which covered most of the hypothalamus, so brain activity estimates could be consistently extracted from each of these subparts in each participant.

**Table 1:** Demographic characteristics of the study sample.

|  | Adults_Morning<br>(AM) | Adults_Evening<br>(AE) | Adolescents<br>(T) | Comparison (t-test) |  |
| --- | --- | --- | --- | --- | --- |
|  |  |  |  | AM vs AE | AE vs T |
| Number of Subjects | 18 | 15 | 16 |  |  |
| Age (mean±SD) | 24.0±2.4 | 24.6±3.8 | 16.8±1.1 | P=0.23 | **P<0.0001** |
| Sex: Female (Male) | 12(6) | 8(7) | 6(10) | P=0.64 | P=0.18 |
| Body mass index<br>(kg/m<sup>2</sup>) | 21.4±2.3 | 22.1±2.0 | 21.8±2.1 | P=0.35 | P=0.69 |
| Depression (BD-II<sup>a</sup>) | 5.6±5.6 | 6.6±3.5 | 6.2±4.8 | P=0.57 | P=0.81 |
| Anxiety Level<br>(BAI<sup>b</sup>) | 5.3±6.5 | 5.5±3.3 | 6.6±7.0 | P=0.89 | P=0.59 |
| Chronotype (HO<sup>c</sup>) | 51.8±7.2 | 43.1±7.3 | 43.4±7.7 | **P=0.002** | P=0.93 |
| Sleep Quality<br>(PSQI<sup>d</sup>) | 3.6±2.9 | 4.1±1.9 | 4.1±1.6 | P=0.62 | P=0.96 |
| Habitual daytime<br>Sleepiness (ESS<sup>e</sup>) | 5.7±3.0 | 6.9±2.4 | 6.0±4.5 | P=0.23 | P=0.52 |
| Season of<br>experiment * | -0.1±0.7 | -0.4±0.5 | -0.4±0.6 | P=0.08 | P=0.70 |
| Seasonality<br>(SPAQ<sup>f</sup>) | 1.0±0.8 | 1.3±0.9 | 0.7±1.0 | P=0.37 | P=0.24 |

<sup>a</sup>. Beck's Depression Inventory II
<sup>b</sup>. Beck's Anxiety Inventory
<sup>c</sup>. Horne and Östberg Questionnaire
<sup>d</sup>. Pittsburg Sleep Quality Index
<sup>e</sup>. EPWORTH Sleepiness Scale
<sup>f</sup>. Seasonal Pattern Assessment Questionnaire
\* Cosine (acquisition day of year\*360/365) : 21 December=0°
\*\* All participants in all groups were right-handed.
Bold values indicate statistical significance at the p<0.05 level.

### Similar Regional Impact of Illuminance on Hypothalamus Activity in the Morning and in the Evening

Our primary analysis regarding time-of-day aimed to detect differences in the overall effects of illuminance changes across the five distinct hypothalamic subparts between the young adults recorded in the morning and those recorded in the evening. For each subpart, an index was computed to reflect the influence of illuminance, derived from the average brain activity estimates that captured how task-related neural activity was modulated by changes in illuminance. As in our previous publication comprising only morning recordings ^9^, the statistical analyses, with the activity of each subpart as the dependent variable, yielded significant differences between the subparts (**p < 0.0001**), indicating that the subparts responded differently to changes in light levels (**Table 1**; **Figure 2-B**). Importantly, however, the analyses did not reveal any time-of-day or time-of-day by hypothalamus subpart interaction (p > 0.59), suggesting that the impact of light was similar in the morning and in the evening (**Figure 2-C**). We further observed a statistical trend for the main effect of task (p = 0.08) and statistically significant main effect of age (within the young adult group; **p = 0.04**), while no significant effects were found for the other covariates.

Again, similar to our previous publication with morning recordings ^9^, the post hoc contrasts revealed that the effect of illuminance variation was consistently greater in the posterior hypothalamus than in the other subparts (**p_corrected_ ≤ 0.0006**), which did not significantly differ from one another (**Table 2**). In addition, as indicated by the absence of a main effect of time-of-day, none of the subparts were significantly different in the morning vs. the evening groups (**Figure 2-C)**. When the impact of illuminance was considered separately in evening and morning groups, post hoc contrasts further indicated that the modulation of the activity of the posterior hypothalamus by the overall changes in illuminance was significantly greater (**p_corrected_ < 0.05**) than that of the other 4 subparts in the morning group, whereas it was greater than that of the inferior-anterior and superior-anterior subparts in the evening group (**Figure 2-C**, **Table 2**). Finally, given the significant difference in chronotype between morning and evening participants, we included chronotype as an additional covariate, but it had no impact on the outcomes of the statistical model (except that the effect of age became a statistical trend, p = 0.07). Chronotype was not significantly associated with hypothalamic subpart activity, neither as a standalone covariate (p = 0.32), nor in interaction with time of day (p = 0.91), hypothalamic subpart (p = 0.16) or their triple interaction (p = 0.63). Overall, our main analyses confirm previous findings that the posterior hypothalamus shows a consistently greater positive modulation by illuminance than other subparts, while this effect does not differ significantly between morning and evening.

**Table 2:**
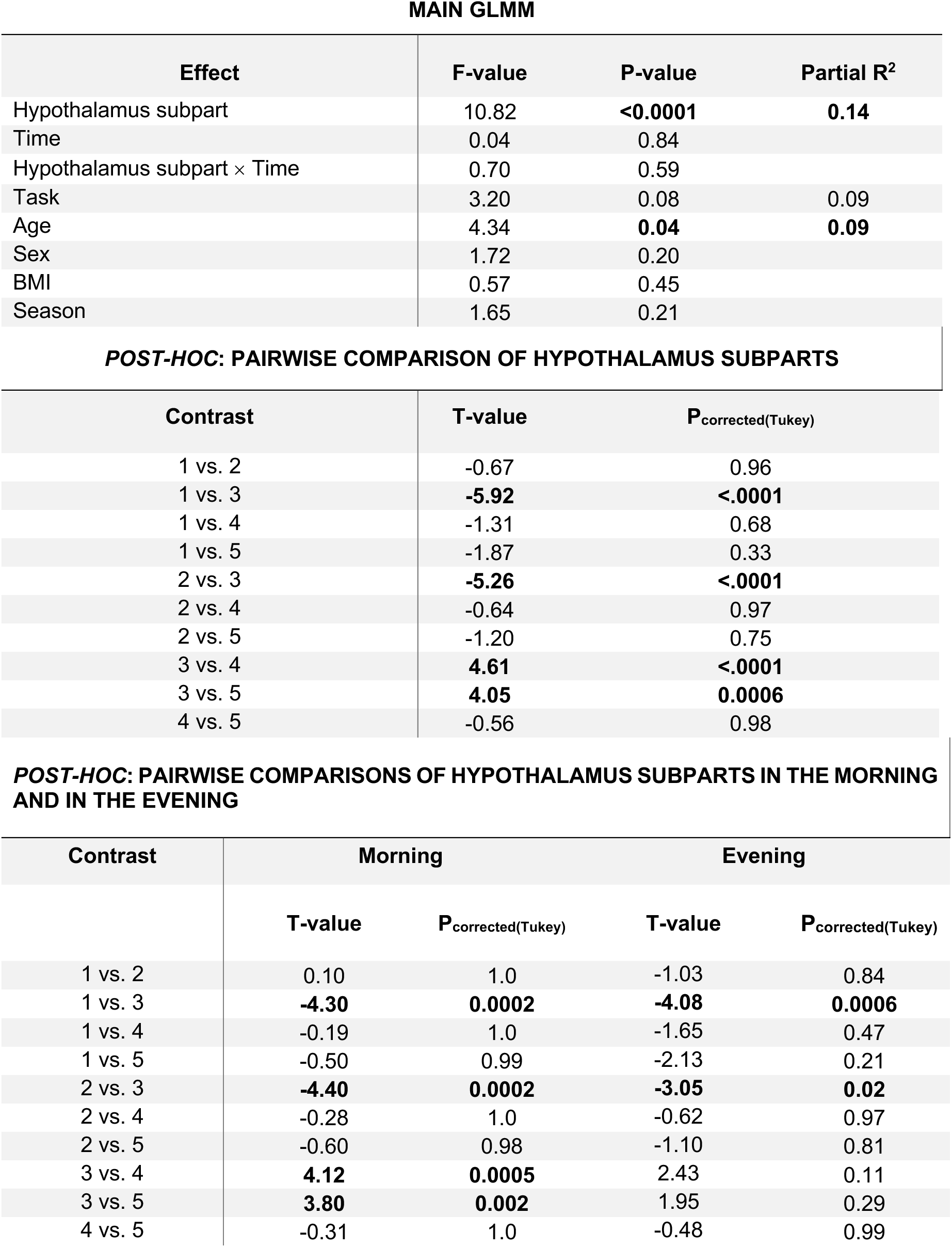
Output of GLMM comparing hypothalamic response in the morning (N=18) vs. evening (N=15) groups with activity of each subpart as the dependent variable, along with outputs of the post hoc test for pairwise comparisons of hypothalamus subparts.

### Opposite Responses to Increased Illuminance in the Posterior and Anterior/Inferior Hypothalamus

To assess whether our main analysis did not overlook local non-linear differences in some subparts, we further investigated the difference in hypothalamic activity across different subparts by examining the responses of each subpart at different illuminance levels. As in our previous publication ^9^, in addition to a main effect of illuminance (**p = 0.0003**), the statistical analysis confirmed that the activity dynamics across illuminance levels differed between the five subparts during the executive task (GLMM; hypothalamus subpart-by-illuminance interaction; **p = 0.0013**) (**Table 3**; **Figure 3**). As in our main analysis, there were no main effects of time-of-day or interaction terms with time-of-day, further supporting that the impact of illuminance on the hypothalamus subparts was similar in the morning and in the evening.

**Figure 3:**
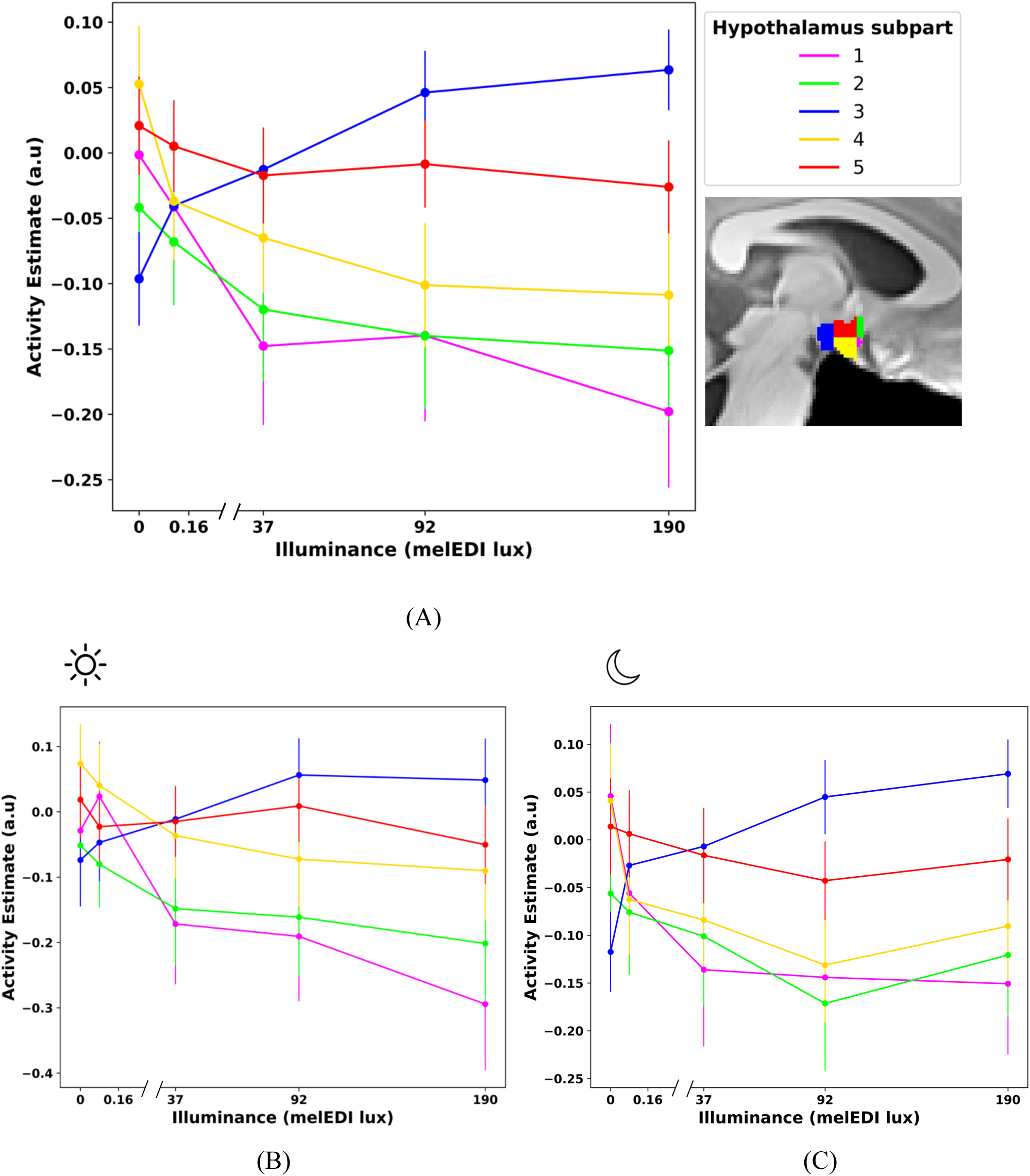
Activity dynamics across illuminance levels for each hypothalamus subpart. Changes of activity estimate (arbitrary unit – a.u.) within each hypothalamus subpart with illuminance in (A) all young adults (N=33), (B) young adult scanned in the morning (N=18) and (C) young adult scanned in the evening (N=15).

**Table 3:** Statistical outputs of GLMM analysis comparing hypothalamic activity in the morning (N=18) vs. evening groups (N=15) in response to each illuminance.

**MAIN GLMM**
| Effect | F-value | P-value | Partial R<sup>2</sup> |
| --- | --- | --- | --- |
| Illuminance | 5.34 | **0.0003** | **0.02** |
| Hypothalamus subpart | 1.32 | 0.26 |  |
| Hypothalamus subpart × Illuminance | 2.42 | **0.0013** | **0.03** |
| Time | 0.74 | 0.40 |  |
| Time × Illuminance | 0.38 | 0.83 |  |
| Time × Hypothalamus subpart | 2.10 | 0.08 | 0.02 |
| Time × Hypothalamus subpart × Illuminance | 0.59 | 0.90 |  |
| Task | 3.46 | 0.06 | 0.01 |
| Age | 7.08 | **0.01** | **0.17** |
| Sex | 0.38 | 0.54 |  |
| BMI | 0.10 | 0.75 |  |
| Season | 3.17 | 0.09 | 0.10 |

Post hoc analyses were conducted to evaluate the impact of illuminance within each subpart (**Suppl. Table S1**). The posterior hypothalamus exhibited increased activity at the two highest illuminance levels (92 and 190 mel-EDI) compared with darkness (**p = 0.016** and **p = 0.005**, respectively). In contrast, the inferior-anterior and superior-anterior hypothalamus exhibited significant decreases in activity (**p < 0.05**) under all blue-enriched conditions (37, 92, and 190 mel-EDI) compared with darkness. Moreover, the inferior-tubular subregion showed reduced activity at the highest illuminance level (190 mel-EDI) compared with the lowest illuminance conditions (0 and 0.16 mel-EDI; **p = 0.01**) as well as decreased activity under the two other blue-enriched light conditions (37 and 92 mel-EDI) compared with darkness (**p = 0.04**). In contrast, the superior-tubular subpart did not show any significant change in activity with varying illuminance levels. When we examined the morning and evening groups separately, we observed similar patterns of activity across both groups: the activity of the posterior hypothalamus increased with increasing illuminance, whereas the activity of the inferior-anterior, superior-anterior, and inferior tubular subparts decreased. In contrast, the superior tubular subpart remained largely unaffected, showing no significant change across the varying illuminance levels.

Overall, the analyses per illuminance level confirm the main findings by demonstrating that hypothalamic subregions respond differently to light intensity. This detailed analysis adds an important layer of insight, revealing that while the posterior hypothalamus, shows increased activation with higher illuminance, others exhibit suppression, and one subpart appears to remain unaffected, reinforcing the distinct impact of light over the hypothalamus subparts.

In both groups, activity increases with illuminance over the posterior hypothalamus (3), decreases over the inferior-anterior (1), superior-anterior (2) and inferior-tubular (4), and remains relatively stable over the superior-tubular (5). Details about the numbers associated to each hypothalamus subpart, can be found in Figure 2-A. The activity estimates of the N0 and N2 tasks are averaged.

### Adolescents Exhibit Different Hypothalamic Response to Illuminance, over the Anterior and Tubular Subparts

Our second primary analysis focused on developmental stage. We aimed to investigate whether the regional response of the hypothalamus to light differed between adolescents and young adults. Given the concerns about evening light exposure in adolescents, data were collected only in the evening, approximately one hour before their habitual bedtime.

As in the first primary analyses, the GLMM yielded a significant main effect of the hypothalamus subpart (**p < 0.0001**) but no significant effect of age group or age group by the hypothalamus subpart interaction (p > 0.5) (**Table 4**). Post hoc contrasts further confirmed that, in young adults, the modulation of the activity of the posterior hypothalamus by the overall changes in illuminance was significantly larger (**p_corrected_ < 0.05**) than that in the other 4 subparts, which did not differ significantly from one another. In adolescents, the modulation of activity in the posterior subpart by illuminance was significantly larger (**p_corrected_ < 0.05**) than that in the inferior anterior and superior anterior subparts (**Figure 4-A**). Overall, this analysis suggests that the linear effect of changes in illuminance is similar across both age groups. However, this does not mean that there are no local non-linear differences in some subparts. To investigate this further, we examined the activity of different hypothalamic subparts at each level of illuminance.

**Figure 4:**
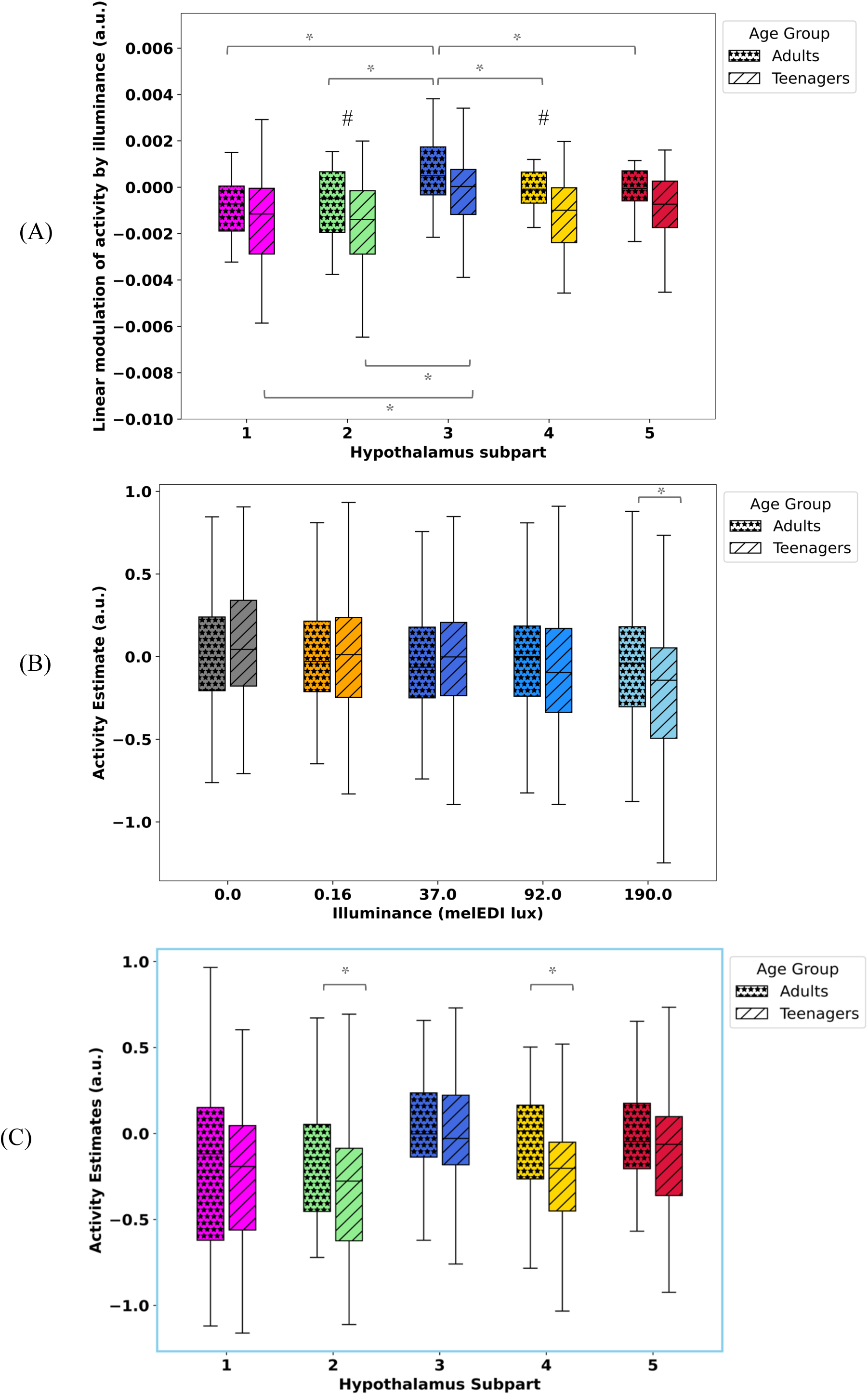
Group comparison of hypothalamic response between adolescents and young adults in the evening. (A) Linear modulation of activity by illuminance (arbitrary unit – a.u.; median (horizontal line), interquartile range (IQR; boxed region), and whiskers extending to Q3 + 1.5 × IQR and Q1 – 1.5 × IQR) of each hypothalamus subpart in response to illuminance variation in adolescent (N=16) and young adult (N=15) groups (main effect of hypothalamus subpart: p < 0.0001). (B) Overall activity estimates (arbitrary unit – a.u.; median (horizontal line), interquartile range (IQR; boxed region), and whiskers extending to Q3 + 1.5 × IQR and Q1 – 1.5 × IQR) of hypothalamus to each illuminance level in adolescents and young adults. There is a significant difference between the groups under the highest illuminance level (190 mel-EDI lux) (P_corrected_ = 0.04) (C) Activity estimates (arbitrary unit – a.u. ; median (horizontal line), interquartile range (IQR; boxed region), and whiskers extending to Q3 + 1.5 × IQR and Q1 – 1.5 × IQR) of each hypothalamus subpart under the under the highest illuminance level (190 mel-EDI lux) in adolescents and young adults. Adolescents show larger deactivation in superior-anterior and inferior-tubular (p = 0.04). The activity estimates of the 0-back and 2-back tasks are averaged (task are presented separately in **Suppl. Figure S3**). Asterisks (*) denote significant differences (p < 0.05).

**Table 4:** Output of GLMM comparing hypothalamic response in adolescents (N=16) vs. young adults (N=15) groups in the evening with activity of each subpart as the dependent variable, along with outputs of the post hoc test for pairwise comparisons of hypothalamus subparts.

| MAIN GLMM |  |  |  |
| --- | --- | --- | --- |
| Effect | F-value | P-value | Partial R<sup>2</sup> |
| Hypothalamus subpart | 10.73 | **<0.0001** | **0.15** |
| Age Group | 0.45 | 0.51 |  |
| Hypothalamus subpart × Age Group | 0.76 | 0.55 |  |
| Task | 0.03 | 0.87 |  |
| Sex | 0.36 | 0.55 |  |
| BMI | 0.54 | 0.47 |  |
| Season | 0.13 | 0.72 |  |

| POST HOC: PAIRWASIE COMPARISON OF HYPOTHALAMUS SUBPARTS IN YOUNG ADULTS AND IN ADOLESCENTS |  |  |  |  |
| --- | --- | --- | --- | --- |
| Contrast | Young Adults |  | Adolescents |  |
|  | T-value | P<sub>corrected(Tukey)</sub> | T-value | P<sub>corrected(Tukey)</sub> |
| 1 vs. 2 | -0.50 | 0.99 | 0.05 | 1.00 |
| 1 vs. 3 | **-5.52** | **<.0001** | **-2.89** | **0.03** |
| 1 vs. 4 | -2.00 | 0.27 | -1.21 | 0.75 |
| 1 vs. 5 | -2.46 | 0.10 | -1.77 | 0.39 |
| 2 vs. 3 | **-4.71** | **<.0001** | **-2.94** | **0.03** |
| 2 vs. 4 | -1.49 | 0.57 | -1.26 | 0.72 |
| 2 vs. 5 | -1.96 | 0.29 | -1.82 | 0.36 |
| 3 vs. 4 | **3.22** | **0.01** | 1.68 | 0.45 |
| 3 vs. 5 | **2.76** | **0.04** | 1.12 | 0.80 |
| 4 vs. 5 | -0.46 | 0.99 | -0.56 | 0.98 |

**Table 5:** Statistical outputs of GLMM analysis comparing hypothalamic activity in adolescents (N=16) vs. young adults (N=15) groups in response to each illuminance in the evening.

| MAIN GLMM |  |  |  |
| --- | --- | --- | --- |
| Effect | F-value | P-value | Partial R<sup>2</sup> |
| Illuminance | 13.35 | <.0001 | 0.04 |
| Hypothalamus subpart | 1.89 | 0.11 |  |
| Hypothalamus subpart × Illuminance | 2.88 | 0.0001 | 0.04 |
| Age group | 0.01 | 0.92 |  |
| Age group × Illuminance | 5.98 | <.0001 | 0.02 |
| Age group × Hypothalamus subpart | 0.81 | 0.52 |  |
| Age group × Hypothalamus subpart × Illuminance | 0.44 | 0.97 |  |
| Task | 6.51 | 0.01 |  |
| Sex | 1.47 | 0.24 |  |
| BMI | 2.74 | 0.11 |  |
| Season | 0.33 | 0.57 |  |
| GROUP COMPARISON BETWEEN OVERALL HYPOTHALAMUS RESPONSE IN ADOLESCENTS AND YOUNG ADULTS AT DIFFERENT ILLUMINANCE LEVELS |  |  |  |
| Illuminance | Contrast | T-value | P<sub>corrected(Tukey)</sub> |
| 0 | Adolescents vs. Adults | 1.27 | 0.20 |
| 0.16 | Adolescents vs. Adults | -0.09 | 0.93 |
| 37 | Adolescents vs. Adults | 0.92 | 0.36 |
| 92 | Adolescents vs. Adults | -0.54 | 0.59 |
| 190 | Adolescents vs. Adults | -1.97 | 0.04 |

The GLMM using the activity of each subpart under each illuminance as the dependent variable, first confirmed distinct changes in activity across subparts under different illuminances (subpart-by-illuminance interaction, **p = 0.0001**), along with the main effects of illuminance (**p < 0.0001**) and task (**p = 0.01**) (**Table 5**). Critically, the GLMM revealed a significant interaction between illuminance and age group (**p < 0.0001**), indicating that changes in activity with variations in illuminance were distinct between adolescents and young adults.

Post hoc analyses indicated that at the highest illuminance level (190 mel-EDI), overall hypothalamic activity differed significantly between the two age groups, with adolescents showing lower activity than young adults did (**p = 0.04**) (**Figure 4-B**, **Table 5**). This difference was largely attributed to the anterior-superior (**p = 0.04**) and inferior-tubular (**p = 0.04**) subparts, which showed larger deactivation in adolescents than in young adults in response to the highest illuminance (**Figure 4-C, Suppl. Table S2**). Considering the activity of each subpart under each illuminance in each group (**Suppl. Table S3, Supp. Figure S2**), we noted that the anterior-inferior subpart significantly decreased in activity with increasing illuminance in both groups. In contrast, the posterior subpart significantly increased only in young adults, whereas the superior-anterior, inferior-tubular and superior-tubular subparts significantly decreased only in teenagers. Therefore, except for the inferior-anterior subpart, 4 subparts of the hypothalamus appeared to display distinct dynamics between age groups, but only between-group differences in the superior-anterior and inferior-tubular subparts reached significance (**Suppl. Table S3**).

### Performance to the Executive Task Is Improved by Light with No Correlation to Hypothalamic Subregion Activity

In the final step, we considered performance to the cognitive task. We focused on the more challenging 2-back condition of the executive task, known to engage higher-order cognitive processes (Collette et al., 2005). Overall task accuracy was high in all participants (mean±SD of accuracy on the 2-back task: 90.0%±7.0% across adults with morning fMRI; 89.6%±7.5% across adults with evening fMRI; and 87.1%±8.2% across adolescents), and performance improved with increasing illuminance, with no impact of time of day or age group (GLMM with all young adults including time of day as covariate while controlling for age, sex, BMI, season and chronotype; main effect of illuminance: F = 3.07, **p = 0.02**, Partial R² = 0.1; **Figure 5-A**; GLMM with adolescents and adults scanned in the evening including age group as covariate while controlling for sex, BMI and season; main effect of illuminance: F = 4.87, **p = 0.001**, Partial R² = 0.2; **Figure 5-B**). Post hoc analyses are overall in line with a performance increase with increasing illuminance (**Table 6**). When investigating whether illuminance-related changes in hypothalamus activity were associated with changes in performance, we found no significant association between the accuracy to the two back task and the activity in any hypothalamic subregion (p > 0.05; **Suppl. Table S4, S5**).

**Figure 5:**
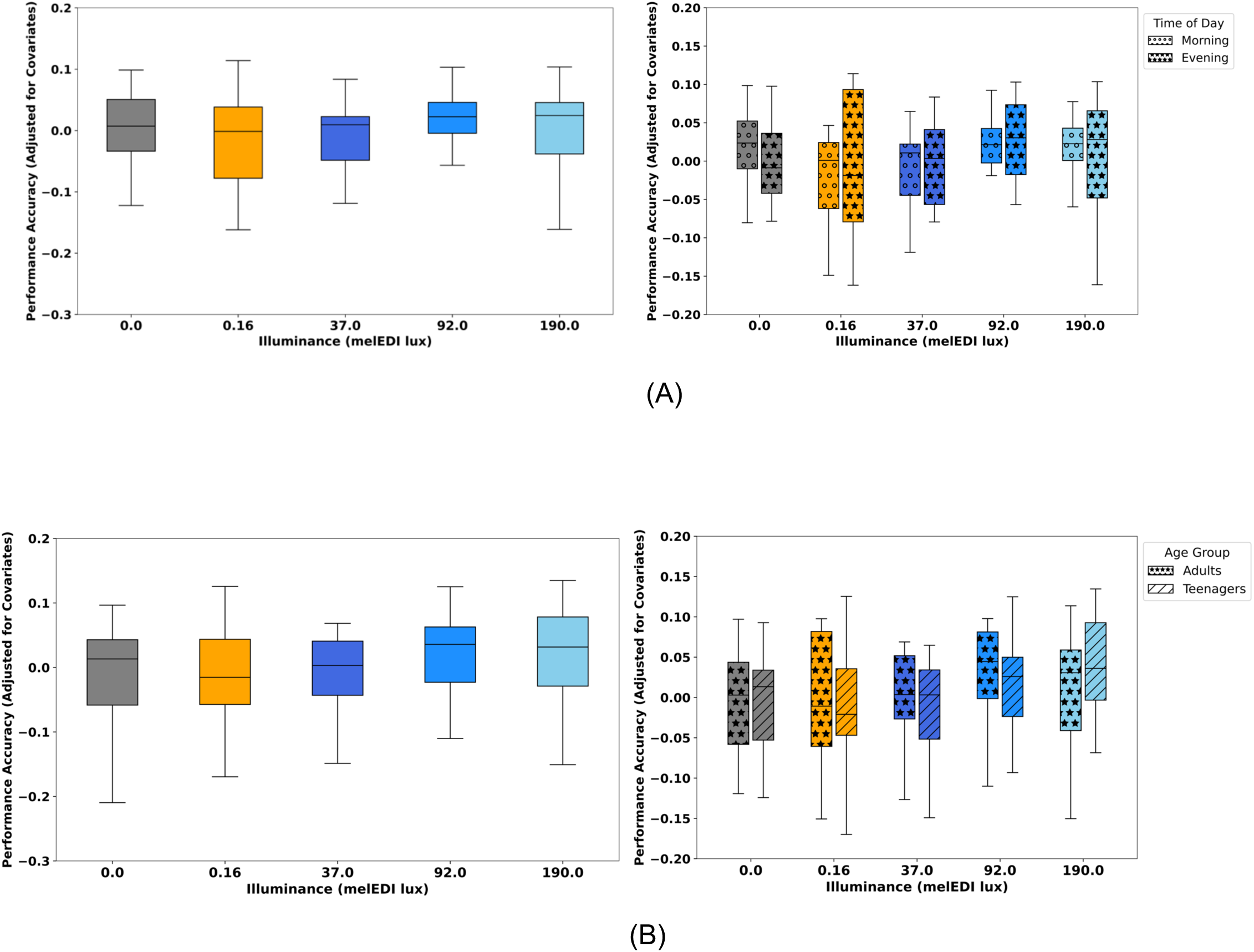
Performance on the 2-back task improved with increasing illuminance, with no significant effect of time of day or age group. (A) accuracy across different illuminance levels in young adults (left: data pooled from both the morning (N=18) and evening (N=15) groups; right: performance in the morning and evening groups separately). (B) accuracy under varying illuminance levels in the evening (left: data pooled from the young adult (N=15) and adolescent (N=16) groups; right: performance in the morning and evening groups separately). In both panels, accuracy values are adjusted for the covariate for display purposes. Box plot showing the median (horizontal line), interquartile range (IQR; boxed region), and whiskers extending to Q3 + 1.5 × IQR and Q1 – 1.5 × IQR.

**Table 6:** Post-hoc analysis: Pairwise comparison of performance under different light levels in adults (pooling data of morning and evening groups) and in the evening (pooling data from adults and adolescents).

| Contrast | ADULTS<br>(MORNING AND EVENING) |  |  | EVENING<br>(ADULTS AND ADOLESCENTS) |  |  |
| --- | --- | --- | --- | --- | --- | --- |
|  | T-value | P<sub>uncorrected</sub> | P<sub>corrected (Tukey)</sub> | T-value | P<sub>uncorrected</sub> | P<sub>corrected (Tukey)</sub> |
| 0 vs. 0.16 | 1.38 | 0.17 | 0.64 | 0.46 | 0.64 | 0.99 |
| 0 vs. 37 | 0.97 | 0.33 | 0.87 | 0.76 | 0.45 | 0.94 |
| 0 vs. 92 | -1.46 | 0.15 | 0.59 | -2.10 | 0.037 | 0.23 |
| 0 vs. 190 | -1.08 | 0.28 | 0.81 | -2.58 | 0.011 | 0.08 |
| 0.16 vs. 37 | -0.41 | 0.68 | 0.99 | 0.28 | 0.78 | 0.99 |
| 0.16 vs. 92 | **-2.84** | **0.005** | **0.04** | -2.56 | 0.011 | 0.08 |
| 0.16 vs. 190 | -2.46 | 0.015 | 0.11 | **-3.06** | **0.003** | **0.02** |
| 37 vs. 92 | -2.43 | 0.016 | 0.11 | **-2.78** | **0.007** | **0.05** |
| 37 vs. 190 | -2.05 | 0.042 | 0.25 | **-3.32** | **0.001** | **0.011** |
| 92 vs. 190 | 0.38 | 0.70 | 0.99 | -0.46 | 0.64 | 0.99 |

## Discussion

The precise mechanism of the biological impact of light on NIF brain function has not been established, particularly in humans, for which translations of findings from animal studies remain scarce. We focused on the hypothalamus, as it is the brain region receiving most of the outputs from ipRGCs and because we previously revealed an opposite response pattern between the anterior and posterior parts of the hypothalamus in response to illuminance variations, when exposed to light in the morning ^9^. Here, we first examined whether the impact of illuminance on the regional activity of hypothalamic was influenced by time-of-day by comparing data collected in the morning and evening in two groups of young adults. We further assess whether illuminance impact varied with developmental stage, by comparing adolescents to young adults in the evening only. Our results reveal that the opposite response to illuminance change of anterior and posterior hypothalamus subparts is similarly present in the morning and in the evening, with the posterior and anterior hypothalamus, respectively, showing increased and decrease activity as illuminance increases. In addition, a detailed analysis of hypothalamus subpart activity revealed distinct dynamics with illuminance change in adolescents vs. young adults, particularly within the superior-anterior subpart and inferior-tubular subpart, which showed a significantly larger decrease in activity as illuminance increased in adolescents. These results suggest that the previously reported time of day differences in the impact of light on brain activity and performance ^12^ may not be primarily driven by prominent regional differences in the hypothalamus. In contrast, previous reports that evening light may affect the circadian system and behavior in adolescents differently ^13^ may be at least in part grounded in regional differences in the response to light across hypothalamus subparts. These findings will contribute to a deeper understanding of the biological effects of light on the brain and offer insights for developing more personalized light interventions.

As in our initial study ^9^ achieving individual hypothalamic nucleus resolution in humans remains unattainable, primarily because of the low contrast between the nuclei, as highlighted in previous works ^34^. We nevertheless employed a standardized, reproducible procedure to parcellate the hypothalamus into consistent subparts ^29^ considered to include (part of) specific nuclei. Yet, this limitation prevents us from attributing our findings to a specific nucleus, and we can suggest only a variety of potential interpretations that would require further investigation. Since we did not observe any time-of-day differences, the interpretations we proposed for the regional impact of illuminance across the hypothalamus in the morning still apply to the evening. We briefly summarize these interpretations in the following lines. The posterior hypothalamus subpart encompasses part of the LH and the TMN, which respectively produce orexin and histamine, both known to promote wakefulness, and animal histology has shown direct projections from ipRGCs to the LH ^2,6,7^. Light may enhance wakefulness by activating these wake-promoting circuits, possibly through an elevated release of orexin and histamine both in the morning and in the evening. This aligns with previous research linking light exposure to improved alertness and reduced subjective sleepiness in the evening ^9^. Orexin is a good candidate for regulating the circadian signal that promotes wakefulness ^35^. If we are indeed facing a change in LH activity with increasing illuminance, this could mean that light similarly induces orexinergic signaling to promote wakefulness both in the morning, shortly after waking, and in the evening, around habitual sleep time.

Similarly, the anterior and tubular subparts of the hypothalamus encompass several nuclei, including two key projection sites of ipRGCs, the SCN and POA, which are involved in circadian and sleep regulation, respectively ^36^, as well as part of the TMN ^3^. The decreased activity we detected in these subparts may reflect a reduction in GABAergic signaling from either of these nuclei ^37–39^, potentially relieving the inhibition of their downstream targets and promoting wakefulness. Regardless of the specific signaling mechanism, the decreased activity we observed in the anterior hypothalamus during the day ^9,10^ appears to be consistently present in the evening. This finding is reminiscent of a previous positron emission tomography (PET) study that reported decreased glucose uptake in the anterior hypothalamus following light exposure at night ^40^. Interestingly, the VLPO has been suggested to deliver inhibitory GABAergic and galaninergic inputs to neurons of the TMN, as well as other components of the ascending monoaminergic arousal system ^41^. Therefore, the opposite activation patterns observed in the anterior region (which includes the VLPO) and the posterior region (which includes the TMN) may be in line with this assumption.

Our findings reveal that the impact of light on hypothalamic activity in young adults extends into the evening as well as the morning, highlighting the importance of carefully managing evening light exposure. The observation of similar effects in the evening raises concerns about the potential disruptive effects of high illuminance levels, especially those exceeding the recommended maximum of 10 mel-EDI lux for evening light ^42^. Such exposure has been shown to disrupt circadian rhythms and sleep ^43^. Broadly speaking, our findings suggest that the hypothalamus may maintain a level of consistency in processing light stimuli across different times of the day. This finding implies that the differential acute biological effects of light on brain function, as reported in previous studies ^12^, may be mediated through other brain regions, such as the pulvinar in the thalamus ^15,44^ or the locus coeruleus in the brainstem.

When comparing adolescents and young adults, only in the evening, we initially found that the overall impact of illuminance on hypothalamic activity did not differ between the two age groups. While this result may not be surprising, given that light likely affects physiology similarly in individuals between the ages of 15 and 30, more detailed analyses revealed differences that were not captured by the linear regression model used to assess the overall impact of illuminance. Specifically, the dynamics of activity changes induced by variations in illuminance appeared to differ across hypothalamic subparts. One notable exception was the inferior-anterior subpart, which displayed qualitatively similar patterns of activity in both age groups, suggesting that the activity of the SCN (or other nuclei within this subpart) and its impact on downstream targets is unchanged in the evening between adolescents and young adults. While we observed some distinct dynamics in the posterior hypothalamic subpart between the two groups, the differences were not statistically significant, so we will not discuss them further. In contrast, the superior-anterior and inferior-tubular subparts of the hypothalamus exhibited a sharper decrease in activity in response to higher light levels, with a more pronounced decrease in adolescents than in young adults.

Compared with young adults, adolescents may therefore exhibit a stronger response to changes in illuminance in the evening, at least in some parts of the hypothalamus. This could contribute to concerns around evening light exposure in this age group, which could be more sensitive to light due to factors such as larger pupil size, clearer lens, and/or a later chronotype ^13,45–48^. This age-related difference in the hypothalamic response might also arise from ongoing neurodevelopmental changes that can potentially alter neurotransmitter dynamics, or the density and connectivity of neural circuits involved in inhibitory responses to light.

If we keep a similar interpretation as for adults in terms of the precise hypothalamus nuclei that may be involved, the age group difference we detect in the evening over the superior-anterior subpart of the hypothalamus could correspond to a stronger response of the POA nuclei and, given their important role in sleep regulation ^6,41^ contribute to sleep disturbances. Similarly, the inferior-tubular subpart could correspond to part of the TMN and imply a reduced GABAergic (or histaminergic) output of the nuclei that could disturb downstream sleep regulation. However, a recent study reported that, while adolescents show greater melatonin suppression by light, their recovery is faster than that of adults ^49^. Specifically, once the light was turned off, the melatonin suppressed by light exposure returned to its pre-exposure level within approximately 50 minutes in adolescents. These findings suggest that although adolescents may experience a stronger acute impact from light exposure, potentially through the specific impacts we detected in the hypothalamus, the effects may dissipate more quickly than they do in younger adults, indicating that young adults may actually be more susceptible to NIF effects.

Similar to our previous findings ^9^, illuminance positively influenced performance on the executive task. However, unlike our earlier study, which had unexpectedly revealed a negative correlation between performance and posterior hypothalamic activity, we found no significant relationship between performance and activity in any hypothalamic subregion in the current study. This discrepancy may be attributed to the different in sample size in the morning or to the environmental (time of day) and individual (age) factors central to the present study and questions the robustness of this earlier observation. As previously ^9^, this reminds that although light-induced performance change is concomitant to light-induced hypothalamus variations, performance, which is an output of cortical activity, may not rely directly and solely on hypothalamus activity.

As with any research, our study bears limitations. The primary limitation is the between-group design, as opposed to a within-subject design, which would have imposed a much higher workload and risk of dropout. Moreover, we have a relatively small sample size within each group, which may have reduced the statistical power of our analyses. This means that there could be weaker effects in our data that we were unable to detect. Additionally, data collection for adolescents was not conducted in the morning because of concerns about school schedule dropout rates, which may have impacted our ability to capture the full range of potential effects. Hence, whether time of day affects regional hypothalamus activity in adolescents remains an open question. We discarded color differences between the light conditions and only considered illuminance as indexed by mel-EDI lux. This does not, for instance, allow us to attribute the findings to a particular photoreceptor class and also means that part of our finding may be attributed to differences in colour. Furthermore, the 5-min exposure to a bright polychromatic light included upon participant arrival to standardize recent prior light history across all groups of participants most likely suppressed melatonin secretion in the evening. Although we cannot exclude it, we consider it is unlike to have greatly contributed to our findings as this short bright light exposure was followed by ∼60 min in dim light (taking into account participants installation and fMRI recording preparation), during which melatonin most likely returned to normal levels ^49,50^. Finally, we used short light exposures, which may have produced different results than longer light exposures did, especially in the relationship between performance and hypothalamus activity.

## Conclusions

In conclusion, this study contributes to the growing body of literature on the relationship between environmental light exposure and an important brain structure, i.e., the hypothalamus, and provides important insights into the differential effects of illuminance on hypothalamic activity across different subparts, times of day, and developmental stages. The posterior hypothalamus seems to be a key region in mediating the arousing effects of light, whereas the anterior hypothalamus may contribute to the stimulating effects of light by inhibiting sleep-promoting circuits, especially in the evening. In the evening, we observed a stronger response to light in adolescents, particularly in subparts of the hypothalamus involved in sleep regulation. This may reinforce concerns about the disruptive effects of evening light on sleep in this age group and highlights the need for careful management of light exposure during key developmental periods. These findings may have important implications for optimizing light exposure to regulate sleep and wakefulness, as well as for improving lighting environments in settings such as schools, workplaces, and therapeutic spaces.

## Declarations Consent for publication

Not applicable

## Availability of data and materials

The processed data and analysis scripts supporting the results included in this manuscript are publicly available via the following open repository: https://gitlab.uliege.be/CyclotronResearchCentre/Public/xxxx (the repository will be created following acceptance/prior to publication of the paper). The raw data could be identified and linked to a single subject and represent a large amount of data. Researchers willing to access the raw data should send a request to the corresponding author (GV). Data sharing will require evaluation of the request by the local Research Ethics Board and the signature of a data transfer agreement (DTA).

## Author Contributions Statement

R.S. I.C, and G.V. designed the research. R.S., F.B., I.C., I.P., and E.B. acquired the data. N.M., F.C., C.P., P.T., L.L. and M.Z. provided valuable insights while acquiring, interpreting, and discussing the data. R.S. analyzed the data which was supervised by G.V. R.S. and G.V. wrote the paper. All authors edited and approved the final version of the manuscript.

## Supporting information

Supplementary materia

## Acknowledgments

The study was conducted within the GIGA-In Vivo Imaging technological platform, ULiège, Belgium. The authors thank Christine Bastin, Alexandre Berger, Christina Schmit, Annick Claes, Christian Degueldre, Catherine Hagelstein, Gregory Hammad, Brigitte Herbillon, Patrick Hawotte, Sophie Laloux, Erik Lambot, Benjamin Lauricella, Pierre Maquet, Eric Salmon and Siya Sherif for their help during the different steps of the study.

