## Supplementary materia for "Regional Hypothalamic Responses to Light Vary Across Developmental Stages but Remain Stable with Time of Day"

### Shared first authorship

~ Joint senior authorship

*Corresponding authors:

Gilles Vandewalle, GIGA-Cyclotron Research Centre-Human Imaging, Bâtiment B30, 8 Allée du Six Août, University of Liège-Sart Tilman, 4000 Liège, Belgium.

Roya Sharifpour, Department of Translational Neuroimaging, Hôpital Erasme, Hôpital Universitaire de Bruxelles, Université libre de Bruxelles, Brussels, Belgium.

**Table S1:** Pair-wise comparison of regional hypothalamic response to different illuminance levels in adults (related to Table 2)

**RESPONSES OF HYPOTHALAMUS SUBPARTS TO EACH ILLUMINANCE**

| Hypothalamus subpart | Contrast | T-value | $P_{corrected(Tukey)}$ |
|---|---|---|---|
| 1 | 0 vs. 0.16 | 0.68 | 0.50 |
| 1 | 0 vs. 37 | **3.06** | **0.002** |
| 1 | 0 vs. 92 | **2.29** | **0.02** |
| 1 | 0 vs. 190 | **4.16** | **<.0001** |
| 1 | 0.16 vs. 37 | **2.38** | **0.02** |
| 1 | 0.16 vs. 92 | 1.61 | 0.11 |
| 1 | 0.16 vs. 190 | **3.48** | **0.0005** |
| 1 | 37 vs. 92 | -0.77 | 0.44 |
| 1 | 37 vs. 190 | 1.10 | 0.27 |
| 1 | 92 vs. 190 | 1.86 | 0.06 |
| 2 | 0 vs. 0.16 | 0.76 | 0.45 |
| 2 | 0 vs. 37 | **2.00** | **0.04** |
| 2 | 0 vs. 92 | **2.46** | **0.01** |
| 2 | 0 vs. 190 | **2.51** | **0.01** |
| 2 | 0.16 vs. 37 | 1.24 | 0.22 |
| 2 | 0.16 vs. 92 | 1.70 | 0.09 |
| 2 | 0.16 vs. 190 | 1.75 | 0.08 |
| 2 | 37 vs. 92 | 0.47 | 0.64 |
| 2 | 37 vs. 190 | 0.52 | 0.61 |
| 2 | 92 vs. 190 | 0.05 | 0.96 |
| 3 | 0 vs. 0.16 | -0.99 | 0.32 |
| 3 | 0 vs. 37 | -1.10 | 0.27 |
| 3 | 0 vs. 92 | **-2.41** | **0.016** |
| 3 | 0 vs. 190 | **-2.79** | **0.005** |
| 3 | 0.16 vs. 37 | -0.10 | 0.92 |
| 3 | 0.16 vs. 92 | -1.42 | 0.16 |
| 3 | 0.16 vs. 190 | -1.80 | 0.07 |
| 3 | 37 vs. 92 | -1.31 | 0.19 |
| 3 | 37 vs. 190 | -1.69 | 0.09 |
| 3 | 92 vs. 190 | -0.38 | 0.70 |
| 4 | 0 vs. 0.16 | 0.74 | 0.46 |
| 4 | 0 vs. 37 | **2.03** | **0.04** |
| 4 | 0 vs. 92 | **2.45** | **0.01** |
| 4 | 0 vs. 190 | **3.31** | **0.001** |
| 4 | 0.16 vs. 37 | 1.30 | 0.19 |
| 4 | 0.16 vs. 92 | 1.72 | 0.09 |

| | | | |
|---|---|---|---|
| 4 | 0.16 vs. 190 | **2.57** | **0.01** |
| 4 | 37 vs. 92 | 0.42 | 0.67 |
| 4 | 37 vs. 190 | 1.27 | 0.20 |
| 4 | 92 vs. 190 | 0.85 | 0.39 |
| 5 | 0 vs. 0.16 | 0.32 | 0.75 |
| 5 | 0 vs. 37 | 1.06 | 0.29 |
| 5 | 0 vs. 92 | 0.76 | 0.44 |
| 5 | 0 vs. 190 | 1.38 | 0.17 |
| 5 | 0.16 vs. 37 | 0.74 | 0.46 |
| 5 | 0.16 vs. 92 | 0.45 | 0.66 |
| 5 | 0.16 vs. 190 | 1.06 | 0.29 |
| 5 | 37 vs. 92 | -0.29 | 0.77 |
| 5 | 37 vs. 190 | 0.32 | 0.75 |
| 5 | 92 vs. 190 | 0.62 | 0.54 |

**Hypothalamus subpart are as follows: inferior-anterior (1), superior-anterior (2), posterior (3), inferior-tubular (4), and superior-tubular. Refer to main text and Figure 2-A for the nuclei included in each subpart.**

**Table S2:** Group comparison of each hypothalamus subpart response at each illuminance level between adolescents and young adults. (related to Table 4)

| Hypothalamus subpart × Illuminance | Contrast | T-value | $P_{corrected(Tukey)}$ |
|---|---|---|---|
| 1 x 0 | Adolescents vs. Adults | 1.02 | 0.31 |
| 1 x 0.16 | Adolescents vs. Adults | -0.93 | 0.35 |
| 1 x 37 | Adolescents vs. Adults | 1.20 | 0.23 |
| 1 x 92 | Adolescents vs. Adults | 0.47 | 0.64 |
| 1 x190 | Adolescents vs. Adults | -0.92 | 0.36 |
| 2 x 0 | Adolescents vs. Adults | 1.00 | 0.32 |
| 2 x 0.16 | Adolescents vs. Adults | -0.45 | 0.65 |
| 2 x 37 | Adolescents vs. Adults | 0.17 | 0.87 |
| 2 x 92 | Adolescents vs. Adults | -1.07 | 0.29 |
| 2 x 190 | Adolescents vs. Adults | -2.01 | **0.04** |
| 3 x 0 | Adolescents vs. Adults | 1.18 | 0.24 |
| 3 x 0.16 | Adolescents vs. Adults | 1.21 | 0.23 |
| 3 x 37 | Adolescents vs. Adults | 1.20 | 0.23 |
| 3 x 92 | Adolescents vs. Adults | 0.03 | 0.97 |
| 3 x 190 | Adolescents vs. Adults | -0.28 | 0.78 |
| 4 x 0 | Adolescents vs. Adults | -0.16 | 0.87 |
| 4 x 0.16 | Adolescents vs. Adults | -0.42 | 0.68 |
| 4 x 37 | Adolescents vs. Adults | -0.14 | 0.89 |
| 4 x 92 | Adolescents vs. Adults | -0.46 | 0.64 |
| 4 x 190 | Adolescents vs. Adults | -2.06 | **0.04** |
| 5 x 0 | Adolescents vs. Adults | 0.90 | 0.37 |
| 5 x 0.16 | Adolescents vs. Adults | 0.31 | 0.75 |
| 5 x 37 | Adolescents vs. Adults | 0.42 | 0.68 |
| 5 x 92 | Adolescents vs. Adults | -0.65 | 0.51 |
| 5 x 190 | Adolescents vs. Adults | -0.86 | 0.39 |

**Hypothalamus subpart are as follows: inferior-anterior (1), superior-anterior (2), posterior (3), inferior-tubular (4), and superior-tubular. Refer to main text and Figure 2-A for the nuclei included in each subpart.**

**Table S3:** Group comparison of between each hypothalamus subpart response to different illuminance levels in adolescents and young adults (related to Table 4)

| Hypothalamus subpart × Age group | Contrast | T-value | $P_{corrected(Tukey)}$ |
|---|---|---|---|
| 1 x Adults | 0 vs. 0.16 | -0.78 | 0.43 |
| 1 x Adults | 0 vs. 37 | 1.97 | **0.05** |
| 1 x Adults | 0 vs. 92 | 2.27 | **0.02** |
| 1 x Adults | 0 vs. 190 | 2.93 | **0.003** |
| 1 x Adults | 0.16 vs. 37 | 2.76 | **0.006** |
| 1 x Adults | 0.16 vs. 92 | 3.05 | **0.002** |
| 1 x Adults | 0.16 vs. 190 | 3.71 | **0.0002** |
| 1 x Adults | 37 vs. 92 | 0.30 | 0.77 |
| 1 x Adults | 37 vs. 190 | 0.95 | 0.34 |
| 1 x Adults | 92 vs. 190 | 0.66 | 0.51 |
| 1 x Adolescents | 0 vs. 0.16 | 1.91 | 0.06 |
| 1 x Adolescents | 0 vs. 37 | 1.78 | 0.07 |
| 1 x Adolescents | 0 vs. 92 | 3.11 | **0.002** |
| 1 x Adolescents | 0 vs. 190 | 5.72 | **<.0001** |
| 1 x Adolescents | 0.16 vs. 37 | -0.13 | 0.90 |
| 1 x Adolescents | 0.16 vs. 92 | 1.20 | 0.23 |
| 1 x Adolescents | 0.16 vs. 190 | 3.81 | **0.0001** |
| 1 x Adolescents | 37 vs. 92 | 1.33 | 0.18 |
| 1 x Adolescents | 37 vs. 190 | 3.94 | **<.0001** |
| 1 x Adolescents | 92 vs. 190 | 2.61 | **0.009** |
| 2 x Adults | 0 vs. 0.16 | -0.11 | 0.91 |
| 2 x Adults | 0 vs. 37 | 0.94 | 0.35 |
| 2 x Adults | 0 vs. 92 | 0.75 | 0.45 |
| 2 x Adults | 0 vs. 190 | 1.10 | 0.27 |
| 2 x Adults | 0.16 vs. 37 | 1.05 | 0.29 |
| 2 x Adults | 0.16 vs. 92 | 0.86 | 0.38 |
| 2 x Adults | 0.16 vs. 190 | 1.21 | 0.23 |
| 2 x Adults | 37 vs. 92 | -0.19 | 0.85 |
| 2 x Adults | 37 vs. 190 | 0.15 | 0.88 |
| 2 x Adults | 92 vs. 190 | 0.34 | 0.73 |
| 2 x Adolescents | 0 vs. 0.16 | 1.91 | 0.06 |
| 2 x Adolescents | 0 vs. 37 | 2.13 | **0.03** |
| 2 x Adolescents | 0 vs. 92 | 3.65 | **0.003** |
| 2 x Adolescents | 0 vs. 190 | 5.33 | **<.0001** |
| 2 x Adolescents | 0.16 vs. 37 | 0.22 | 0.82 |
| 2 x Adolescents | 0.16 vs. 92 | 1.75 | 0.08 |
| 2 x Adolescents | 0.16 vs. 190 | 3.42 | **0.0006** |

| | | | |
|---|---|---|---|
| 2 x Adolescents | 37 vs. 92 | 1.53 | 0.13 |
| 2 x Adolescents | 37 vs. 190 | 3.20 | **0.001** |
| 2 x Adolescents | 92 vs. 190 | 1.67 | 0.09 |
| 3 x Adults | 0 vs. 0.16 | -0.33 | 0.74 |
| 3 x Adults | 0 vs. 37 | -1.15 | 0.25 |
| 3 x Adults | 0 vs. 92 | -1.95 | **0.05** |
| 3 x Adults | 0 vs. 190 | -2.04 | **0.04** |
| 3 x Adults | 0.16 vs. 37 | -0.81 | 0.42 |
| 3 x Adults | 0.16 vs. 92 | -1.62 | 0.10 |
| 3 x Adults | 0.16 vs. 190 | -1.70 | 0.09 |
| 3 x Adults | 37 vs. 92 | -0.81 | 0.42 |
| 3 x Adults | 37 vs. 190 | -0.89 | 0.37 |
| 3 x Adults | 92 vs. 190 | -0.08 | 0.94 |
| 3 x Adolescents | 0 vs. 0.16 | -0.39 | 0.70 |
| 3 x Adolescents | 0 vs. 37 | -1.23 | 0.22 |
| 3 x Adolescents | 0 vs. 92 | -0.42 | 0.67 |
| 3 x Adolescents | 0 vs. 190 | -0.08 | 0.94 |
| 3 x Adolescents | 0.16 vs. 37 | -0.84 | 0.40 |
| 3 x Adolescents | 0.16 vs. 92 | -0.04 | 0.97 |
| 3 x Adolescents | 0.16 vs. 190 | 0.31 | 0.76 |
| 3 x Adolescents | 37 vs. 92 | 0.80 | 0.42 |
| 3 x Adolescents | 37 vs. 190 | 1.15 | 0.25 |
| 3 x Adolescents | 92 vs. 190 | 0.34 | 0.73 |
| 4 x Adults | 0 vs. 0.16 | 0.34 | 0.73 |
| 4 x Adults | 0 vs. 37 | 1.15 | 0.25 |
| 4 x Adults | 0 vs. 92 | 1.51 | 0.13 |
| 4 x Adults | 0 vs. 190 | 1.446 | 0.15 |
| 4 x Adults | 0.16 vs. 37 | 0.81 | 0.42 |
| 4 x Adults | 0.16 vs. 92 | 1.17 | 0.24 |
| 4 x Adults | 0.16 vs. 190 | 1.12 | 0.26 |
| 4 x Adults | 37 vs. 92 | 0.36 | 0.72 |
| 4 x Adults | 37 vs. 190 | 0.30 | 0.76 |
| 4 x Adults | 92 vs. 190 | -0.06 | 0.95 |
| 4 x Adolescents | 0 vs. 0.16 | 0.71 | 0.48 |
| 4 x Adolescents | 0 vs. 37 | 1.16 | 0.25 |
| 4 x Adolescents | 0 vs. 92 | 1.98 | **0.05** |
| 4 x Adolescents | 0 vs. 190 | 4.15 | **<.0001** |
| 4 x Adolescents | 0.16 vs. 37 | 0.45 | 0.65 |
| 4 x Adolescents | 0.16 vs. 92 | 1.27 | 0.20 |
| 4 x Adolescents | 0.16 vs. 190 | 3.44 | **0.0006** |
| 4 x Adolescents | 37 vs. 92 | 0.82 | 0.41 |
| 4 x Adolescents | 37 vs. 190 | 2.99 | **0.003** |
| 4 x Adolescents | 92 vs. 190 | 2.16 | **0.03** |

| | | | |
|---|---|---|---|
| 5 x Adults | 0 vs. 0.16 | 0.39 | 0.70 |
| 5 x Adults | 0 vs. 37 | 0.30 | 0.77 |
| 5 x Adults | 0 vs. 92 | -0.33 | 0.74 |
| 5 x Adults | 0 vs. 190 | 0.37 | 0.71 |
| 5 x Adults | 0.16 vs. 37 | -0.09 | 0.93 |
| 5 x Adults | 0.16 vs. 92 | -0.73 | 0.47 |
| 5 x Adults | 0.16 vs. 190 | 0.02 | 0.98 |
| 5 x Adults | 37 vs. 92 | -0.63 | 0.53 |
| 5 x Adults | 37 vs. 190 | 0.07 | 0.94 |
| 5 x Adults | 92 vs. 190 | 0.70 | 0.48 |
| 5 x Adolescents | 0 vs. 0.16 | 1.23 | 0.22 |
| 5 x Adolescents | 0 vs. 37 | 0.99 | 0.32 |
| 5 x Adolescents | 0 vs. 92 | 1.83 | 0.07 |
| 5 x Adolescents | 0 vs. 190 | 2.84 | **0.005** |
| 5 x Adolescents | 0.16 vs. 37 | -0.24 | 0.81 |
| 5 x Adolescents | 0.16 vs. 92 | 0.60 | 0.55 |
| 5 x Adolescents | 0.16 vs. 190 | 1.61 | 0.11 |
| 5 x Adolescents | 37 vs. 92 | 0.84 | 0.40 |
| 5 x Adolescents | 37 vs. 190 | 1.85 | 0.06 |
| 5 x Adolescents | 92 vs. 190 | 1.01 | 0.31 |

**Hypothalamus subpart are as follows: inferior-anterior (1), superior-anterior (2), posterior (3), and inferior-tubular (4), superior-tubular. Refer to main text and figure 1 for the nuclei included in each subpart.**

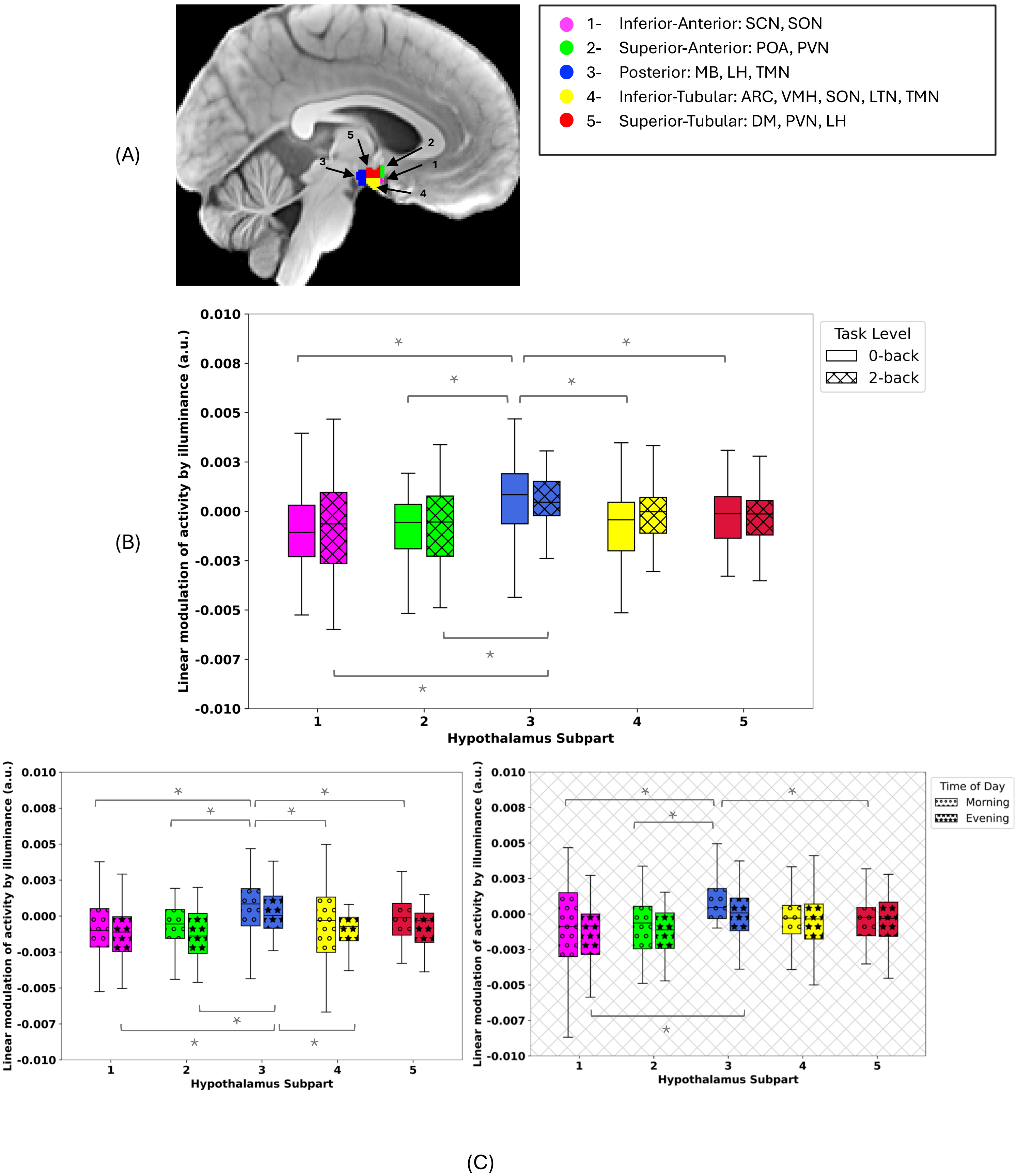

(A)
5
2
3
1
4
1- Inferior-Anterior: SCN, SON
2- Superior-Anterior: POA, PVN
3- Posterior: MB, LH, TMN
4- Inferior-Tubular: ARC, VMH, SON, LTN, TMN
5- Superior-Tubular: DM, PVN, LH
(B)
Linear modulation of activity by illuminance (a.u.)
Hypothalamus Subpart
Task Level
0-back
2-back
0.010
0.008
0.005
0.003
0.000
-0.003
-0.005
-0.007
-0.010
1
2
3
4
5
Time of Day
Morning
Evening
(C)

**Figure S1: Regional hypothalamus response to light in young adults in the morning and evening groups at each task level: 0-back, 2-back**

**(A)** Hypothalamus parcellation into five subparts encompassing the nuclei mentioned in the right inset. ARC: arcuate nucleus; DMH: dorsomedial nucleus; LH: lateral hypothalamus; LTN: lateral tubular nucleus; MB: mamillary body; POA: preoptic area; PVN: paraventricular nucleus; PNH: posterior nucleus of the hypothalamus; SCN: suprachiasmatic nucleus; SON: supraoptic nucleus; TMN: tuberomammillary nucleus; VMN: ventromedial nucleus.

**(B)** Linear modulation of activity by illuminance (arbitrary unit – a.u.; median (horizontal line), interquartile range (IQR; boxed region), and whiskers extending to Q3 + 1.5 × IQR and Q1 – 1.5 × IQR.)) of each hypothalamus subpart in response to illuminance variation during each task level (Morning and evening subject together; N=33) showing increased impact of light in subpart three (posterior) relative to all the other subparts in the morning and the anterior parts in the evening (main effect of hypothalamus subpart: $p<0.0001$).

**(C)** Group comparison of linear modulation of activity by illuminance (a.u.; median (horizontal line), interquartile range (IQR; boxed region), and whiskers extending to Q3 + 1.5 × IQR and Q1 – 1.5 × IQR.)) of each hypothalamus subpart in response to illuminance variation in the morning (N=18) vs. in the evening (N=15) during 0-back (left) and 2-back (right) (main effect of time of day: $p= 0.29$; interaction of hypothalamus subpart and time of day: $p = 0.62$; interaction of hypothalamus subpart and task level: $p = 0.58$; time of day and task level: $p = 0.73$; interaction of hypothalamus subpart, time of day and task level: $p = 0.95$).

Asterisks (*) denote significant differences ($p<0.05$).

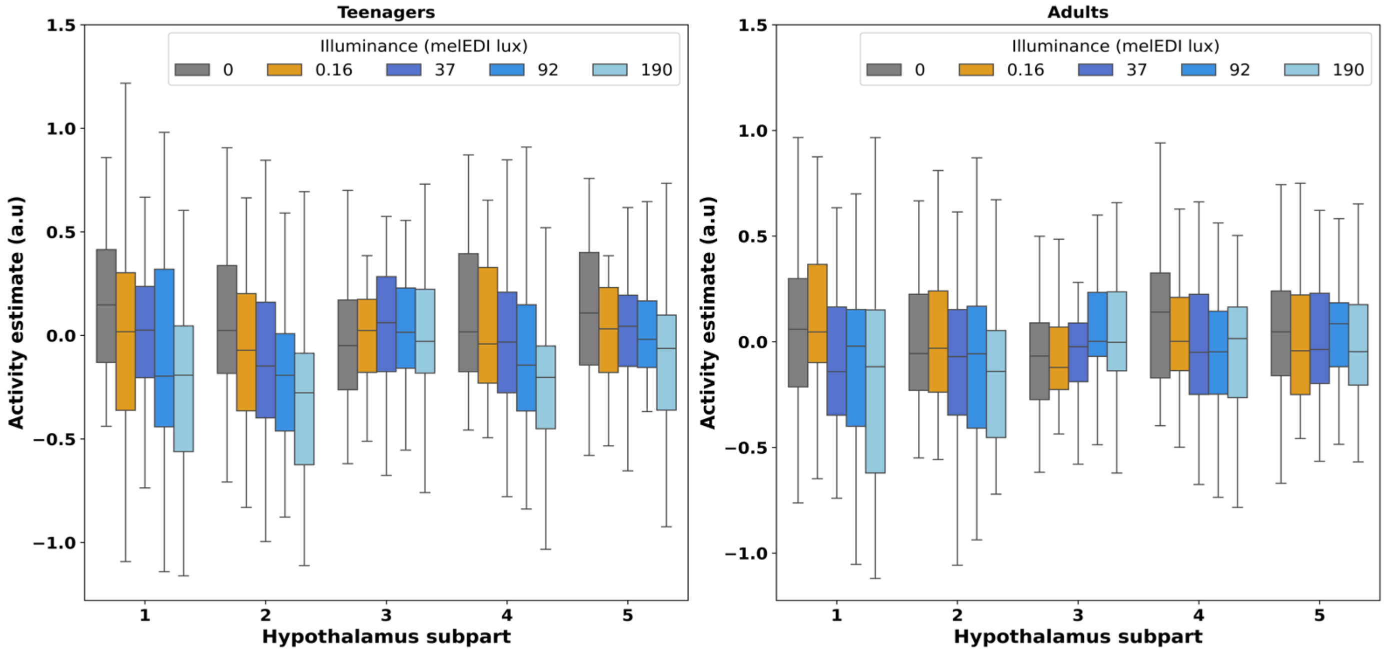


xx

**Figure S2: Response of each hypothalamus subpart to each illuminance level** in adolescents (left) and young adults (right). Refer to **Table S3** for statistical outcomes.

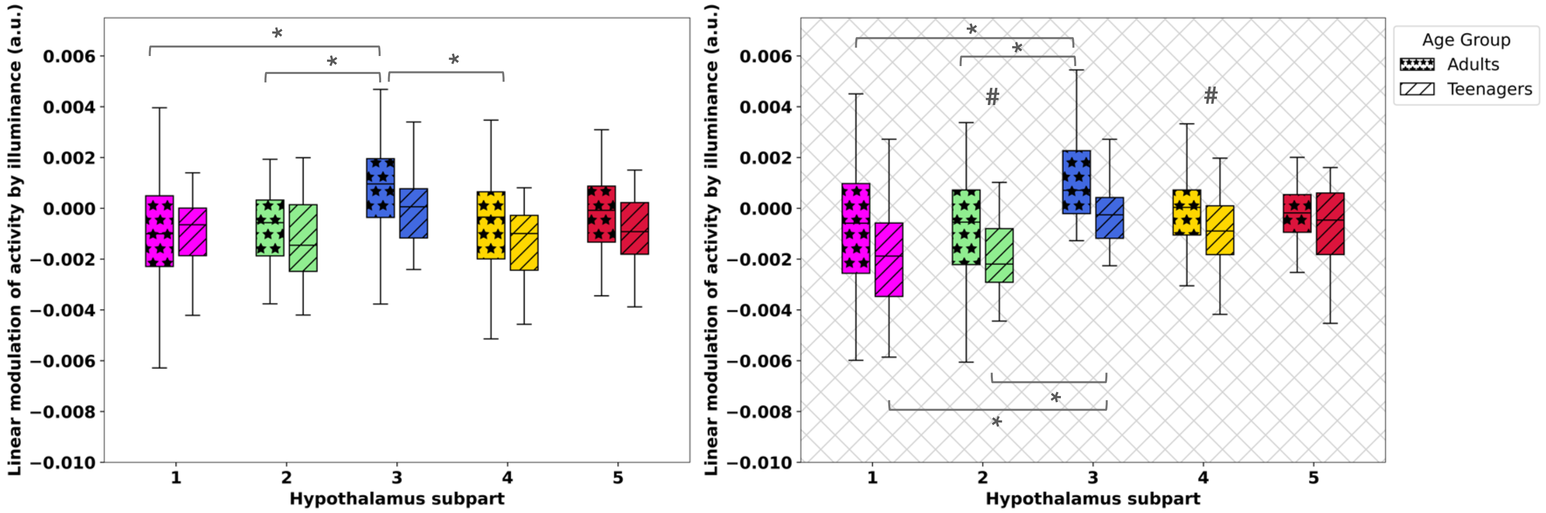
Linear modulation of activity by illuminance (a.u.)
0.006
0.004
0.002
0.000
−0.002
−0.004
−0.006
−0.008
−0.010
1
2
3
4
5
Hypothalamus subpart
Age Group
Adults
Teenagers

(A)

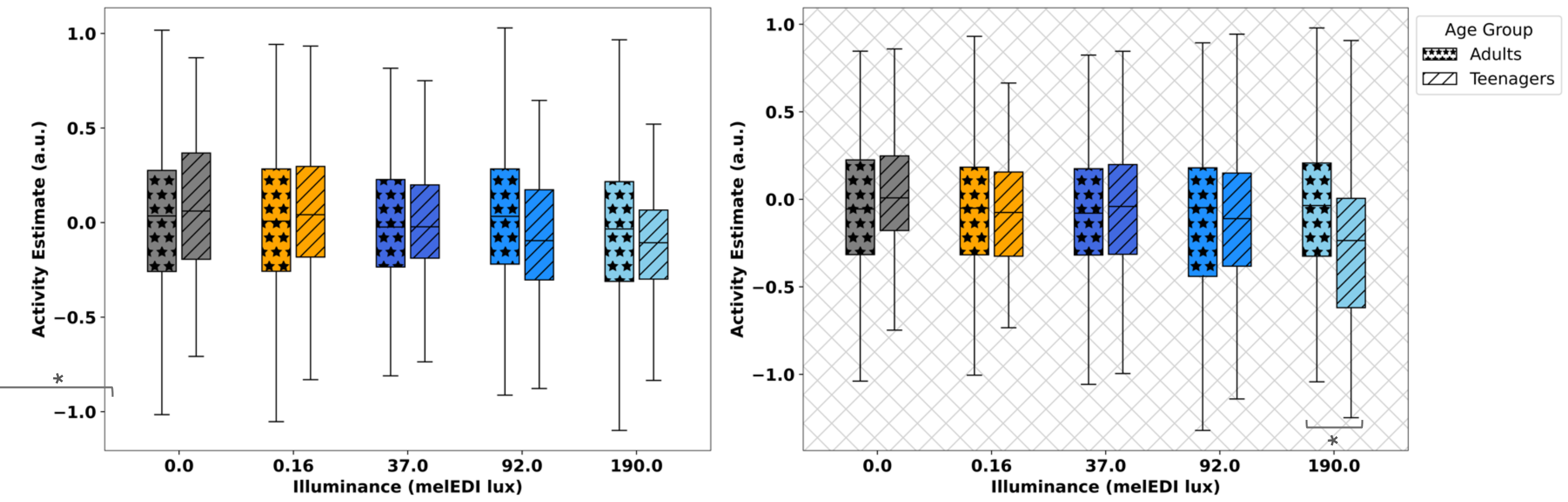
Activity Estimate (a.u.)
1.0
0.5
0.0
−0.5
−1.0
0.0
0.16
37.0
92.0
190.0
Illuminance (melEDI lux)
Age Group
Adults
Teenagers

(B)

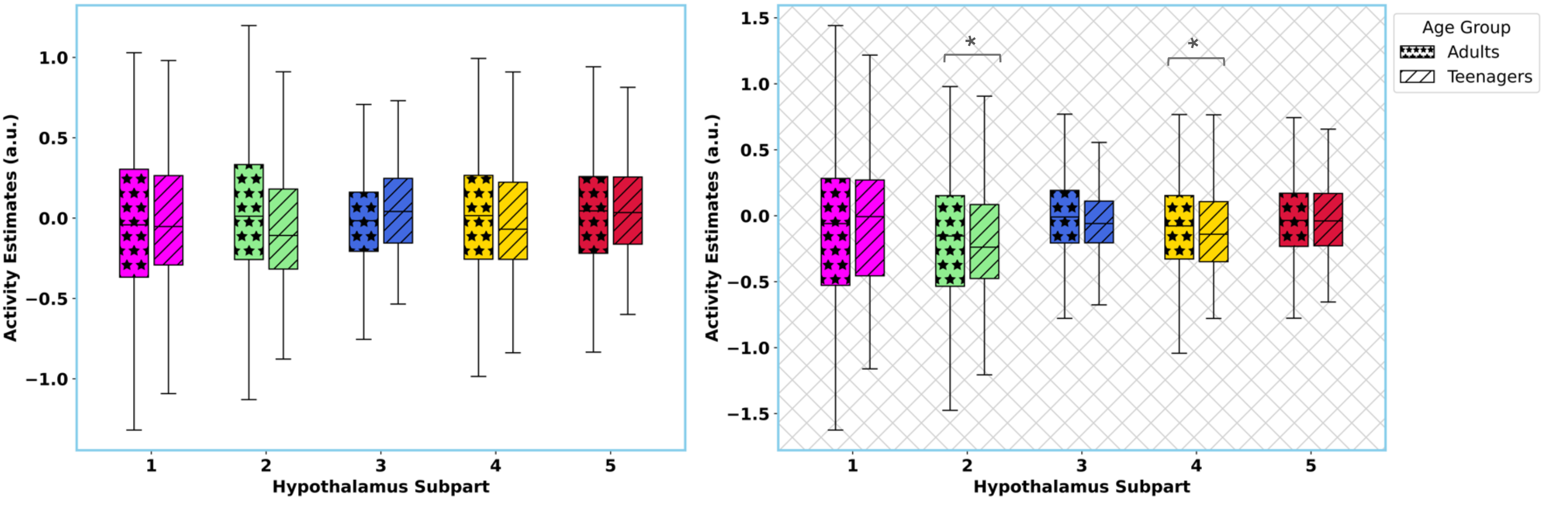
Activity Estimates (a.u.)
1.5
1.0
0.5
0.0
−0.5
−1.0
−1.5
1
2
3
4
5
Hypothalamus Subpart
Age Group
Adults
Teenagers

(C)

**Figure S3: Group comparison of hypothalamic response between adolescents and young adults in the evening at each task level: 0-back and 2-back.**

**(A)** Linear modulation of activity by illuminance (arbitrary unit – a.u.; median (horizontal line), interquartile range (IQR; boxed region), and whiskers extending to Q3 + 1.5 × IQR and Q1 – 1.5 × IQR) of each hypothalamus subpart in response to illuminance variation in adolescent (N=16) and young adult (N=15) groups, during 0-back (left) and 2-back (right).

**(B)** Overall Activity estimates (arbitrary unit – a.u.; median (horizontal line), interquartile range (IQR; boxed region), and whiskers extending to Q3 + 1.5 × IQR and Q1 – 1.5 × IQR) of hypothalamus to each illuminance level in adolescents and young adults. There is a significant difference between the groups under the highest illuminance level (190 mel-EDI lux), during 0-back (left) and 2-back (right).

**(C)** Activity estimates (arbitrary unit – a.u.; median (horizontal line), interquartile range (IQR; boxed region), and whiskers extending to Q3 + 1.5 × IQR and Q1 – 1.5 × IQR) of each hypothalamus subpart under the highest illuminance level (190 mel-EDI lux) in adolescents and young adults. Adolescents show larger deactivation in superior-anterior and inferior-tubular, during 0-back (left) and 2-back (right).

Asterisks (*) denote significant differences ($p<0.05$).

Refer to **Table 5 in the main text** for statistical outcomes.

**Table S4:** Output of GLMM comparing adults' task performance in the morning (N=18) vs. evening (N=15) with task accuracy as the dependent variable.

**GLMM**

| Effect | F-value | P-value | Partial $R^2$ |
|---|---|---|---|
| Activity estimate | 0.00 | 1.00 | - |
| Hypothalamus subpart | 0.00 | 1.00 | - |
| Hypothalamus subpart × Activity estimate | 0.00 | 1.00 | - |
| Time of Day | 11.38 | 0.22 | - |
| Illuminance | 0.23 | 0.71 | - |
| Age | 25.60 | 0.12 | - |
| Sex | 21.40 | 0.13 | - |
| BMI | 29.60 | 0.12 | - |
| Season | 0.75 | 0.55 | - |
| Chronotype | 2.74 | 0.35 | - |

* Considering each subpart one by one did not lead to significant association (p>0.05)

**Table S5:** Output of GLMM comparing evening task performance in adults (N=15) vs. adolescents (N=16) with task accuracy as the dependent variable.

**GLMM**

| Effect | F-value | P-value | Partial $R^2$ |
|---|---|---|---|
| Activity estimate | 0.00 | 1.00 | - |
| Hypothalamus subpart | 0.00 | 1.00 | - |
| Hypothalamus subpart × Activity estimate | 0.00 | 1.00 | - |
| Age Group | 0.20 | 0.65 | - |
| Illuminance | 12.01 | 0.21 | - |
| Sex | 12.84 | 0.0004 | - |
| BMI | 2.24 | 0.13 | - |
| Season | 15.98 | **<.0001** | **0.02** |

* Considering each subpart one by one did not lead to significant association (p>0.05)
